# Movement-Dependent Electrical Stimulation for Volitional Strengthening of Cortical Connections in Behaving Monkeys

**DOI:** 10.1101/2021.09.03.458906

**Authors:** Samira Moorjani, Sarita Walvekar, Eberhard E. Fetz, Steve I. Perlmutter

## Abstract

Correlated activity of neurons can lead to long-term strengthening or weakening of the connections between them. In addition, the behavioral context, imparted by execution of physical movements or the presence of a reward, can modulate the plasticity induced by Hebbian mechanisms. In the present study, we have combined behavior and induced neuronal correlations to strengthen connections in the motor cortex of adult behaving monkeys. Correlated activity was induced using an electrical-conditioning protocol in which stimuli gated by voluntary movements were used to produce co-activation of neurons at motor-cortical sites involved in those movements. Delivery of movement-dependent stimulation resulted in small increases in the strength of associated cortical connections immediately after conditioning. Remarkably, when paired with further repetition of the movements that gated the conditioning stimuli, there were substantially larger gains in the strength of cortical connections, that occurred in a use-dependent manner, without delivery of additional conditioning stimulation. In the absence of such movements, little change was observed in the strength of motor-cortical connections. Performance of the motor behavior in the absence of conditioning also did not produce any changes in connectivity. Our results show that combining movement-gated stimulation with further natural use of the “conditioned” pathways after stimulation ends can produce use-dependent strengthening of connections in adult primates, highlighting an important role for behavior in cortical plasticity. Our data also provide strong support for combining movement-gated stimulation with use-dependent physical rehabilitation for strengthening connections weakened by a stroke or spinal-cord injury.

**Significance Statement:** We describe an electrical-conditioning protocol in adult behaving monkeys in which stimuli gated by voluntary movements were used to strengthen connections between motor-cortical neurons involved in those movements. Movement-gated stimulation created a plastic landscape in which repetition of the movements that gated conditioning stimuli produced strengthening of cortical connections, in a use-dependent manner, long after stimulation had ended, a finding that is both novel and unique. In the absence of such behavior, little change was observed in the strength of connections. Similarly, movements alone did not produce any changes in connectivity. Our data highlight a critical role for behavior in plasticity and provide strong support for combining movement-gated stimulation with use-dependent rehabilitation for strengthening connections weakened by injury or disease.

## Introduction

Spike-timing-dependent plasticity (STDP) refers to correlated activity of neurons that leads to long-term strengthening or weakening of the connections between them, depending on the order of firing of pre- and postsynaptic cells (1–7). Although much evidence supports the necessity of correlated neuronal activity for synaptic plasticity to occur, physical movements and behavioral factors, such as motivation, stress, attention and reinforcement, which occur at very different timescales from STDP (8), are also known to play crucial roles in modulation of functional plasticity (9–17). This adds a layer of complexity to the neuronal computations underlying plasticity processes and the recovery, mediated by such mechanisms, from an injury to the central nervous system (18, 19). Thorndike argued that a connection is significantly modified only when associated with outcomes important to the animal’s behavior (20, 21), suggesting a volitional dimension to the control of plasticity, perhaps through the release of neuromodulators such as dopamine and acetylcholine that play important roles in reward (22, 23) and attention (24, 25) circuits, respectively. Correlated activity of neurons, while often necessary, is not always sufficient to induce plasticity. However, a relatively small number of studies have explored the role of behavioral context in the regulation of synaptic plasticity (9–14, 17, 19).

Motivated by the Hebb-Stent (or STDP) learning rule (1–5) and Thorndike’s law of effect (20), we sought to determine the extent to which an activity-dependent stimulation protocol that produced co-activation of neurons at two motor-cortical sites could be exploited to modulate the connectivity between them, both in the presence and absence of a relevant behavioral context. While a number of laboratories (summarized in 26), including ours, have developed conditioning paradigms for inducing STDP, the role of behavior in synaptic plasticity remains largely uninvestigated. To dissect this role, we implemented an electrical-conditioning paradigm in adult behaving monkeys in which stimuli were delivered during volitional movements that activated neurons at a presynaptic site. The gated stimuli were delivered to a postsynaptic site in an attempt to boost the firing of postsynaptic neurons while the presynaptic neurons were active. Delivery of movement-gated stimulation resulted in small increases in the strength of cortical connections immediately after conditioning. Remarkably, when paired with further repetition of the movements that gated the conditioning stimuli, there were substantially larger increases in the strength of connections, without additional delivery of conditioning stimulation. Importantly, neither behavior alone nor conditioning alone produced similar effects. Note that the term “behavior,” as used here, involves both voluntary movements and accompanying behavioral modulators, such as stress, motivation, attention and reinforcement. Each of these variables may play distinct roles in modulation of plasticity (9–19), but they were not controlled or differentiated in our study.

Our results suggest that movement-gated stimulation creates a plastic landscape in which repetition of the behavioral context presented during conditioning drives cortical strengthening long after stimulation has ended. Taken together, our data highlight a crucial role for behavior in modulation of synaptic plasticity. They also provide support for combining movement-gated stimulation with use-dependent physical rehabilitation for strengthening motor pathways weakened by injury or disease.

## Results

### Movement-dependent stimulation protocol and experimental design

Two adult monkeys (Y and U) were trained to perform a randomly-alternating center-out wrist flexion-extension target-tracking task with their right hands. Monkeys were trained to hold the cursor inside presented targets for 1.6 seconds and were rewarded with fruit sauce at a variable reinforcement ratio after successful completion of trials. After learning the target-tracking task, monkeys received chronic bilateral implants, consisting of custom-made electrode arrays (whose design is described in 27; also see SI Appendix, Materials and Methods), with platinum-iridium (Pt/Ir) microwires targeting sites in layer V of the sensorimotor cortex. All tested site pairs were in the left primary motor cortex in both animals, contralateral to their responding hands.

Motor outputs of two reciprocally-connected neuronal sites, termed Ntest and Nstim, in the primary motor cortex were assessed using trains of intracortical microstimulation (ICMS; for details, see SI Appendix, Materials and Methods; Figure 1A). Movement-related activity at Ntest and Nstim was inferred from their motor outputs (Mtest and Mstim, respectively). The strength of synaptic connections between cortical sites was documented by the size of evoked potentials (EPs; 28, 29) in local-field-potential (LFP) recordings, referred to as EP amplitude, recorded at one site after biphasic charge-balanced test stimuli were delivered at a second site (Figures 1B and 1D). The perturbation of activity in one set of neurons followed by the quantification of its impact, or the evoked response, at other sites provides a directed approach to assess connectivity within and across brain regions, given a direct correlation between EP amplitudes and the strength of connections (28).

**Figure 1.**
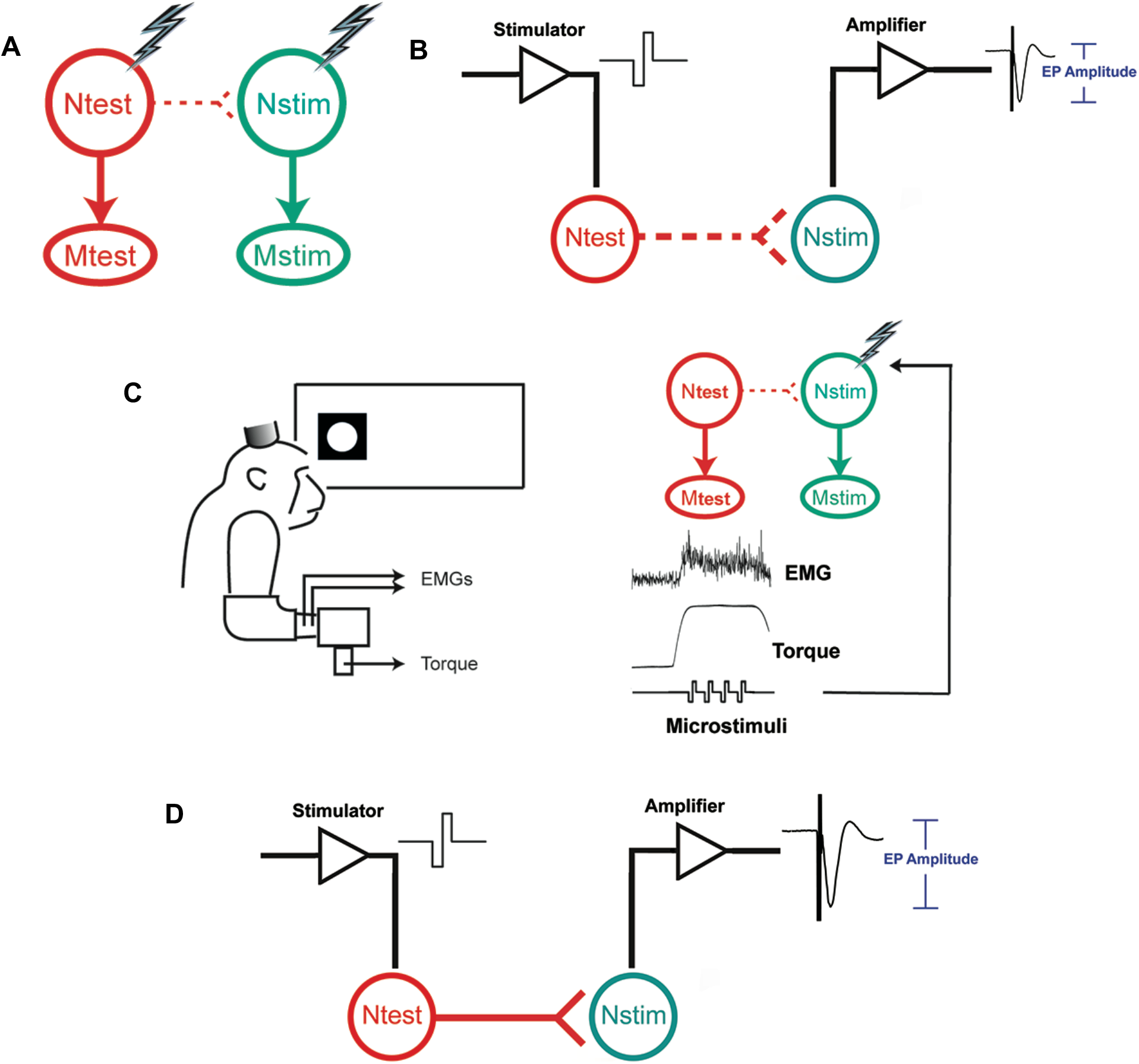
Electrical-conditioning protocol and experimental design. (A) Trains of microstimuli were delivered at two sites in the motor cortex, Ntest and Nstim, to document their motor outputs (Mtest and Mstim, respectively). (C) During conditioning, cursor hold inside the Mtest target gated delivery of conditioning stimuli to Nstim while the monkey performed a wrist flexion-extension target-tracking task. (B, D) Setup for assessing conditioning changes. Cortical responses at Nstim evoked by delivery of test stimuli at Ntest were used to document changes in the strength of the connection from Ntest to Nstim before (B) and after (D) delivery of movement-gated stimulation.

Movement-gated stimulation was delivered to Nstim during movements that were expected to activate neurons at Ntest. The conditioning stimulation likely boosted the firing of neurons at Nstim and produced co-activation of the two cortical sites. To achieve this, delivery of conditioning stimuli at Nstim was gated by cursor hold inside the Mtest target. For example, if the motor output of Ntest was flexion, the monkey received flexion-gated stimulation at Nstim while the cursor was held inside the flexion target, as shown in Figure 1C. Hence, conditioning stimuli were delivered during the plateau phase of torque or the tonic phase of Mtest electromyographic (EMG) activity. Movement-gated stimulation was delivered at 10 Hz at an intensity that was subthreshold for movement as long as the cursor remained in the Mtest target. Delivery of conditioning was preceded and followed by delivery of test stimuli, which were also subthreshold for movement, at Ntest to assess the strength of its connection to Nstim (Figures 1B and 1D). A detailed description of how conditioning and test currents were chosen in our experiments is included in SI Appendix, Materials and Methods.

Thus, two types of electrical stimuli were delivered in our experiments: conditioning stimuli, which were delivered during voluntary movements, and test stimuli, which were used to document motor outputs and the strength of cortical connections. Conditioning stimuli were delivered during wrist flexion or extension; test stimuli were always delivered with the monkey’s wrist at rest.

### Immediate effects of movement-gated conditioning

Figure 2 shows stimulus-triggered averages (StTAs) of cortical field potentials at Nstim evoked by delivery of test stimuli at Ntest. The time course of the experimental session is shown in the inset. Here, movement-gated stimulation was delivered for 90 minutes and was preceded and followed by delivery of test stimuli to document changes in connectivity from Ntest to Nstim. Test stimuli were delivered at several time points post conditioning up to an hour. The monkey performed the flexion-extension target-tracking task throughout the session. Connectivity changes were quantified as percent changes in the average EP amplitude after conditioning relative to the preconditioning level, according to the following equation.

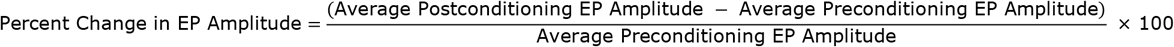

**Figure 2.**
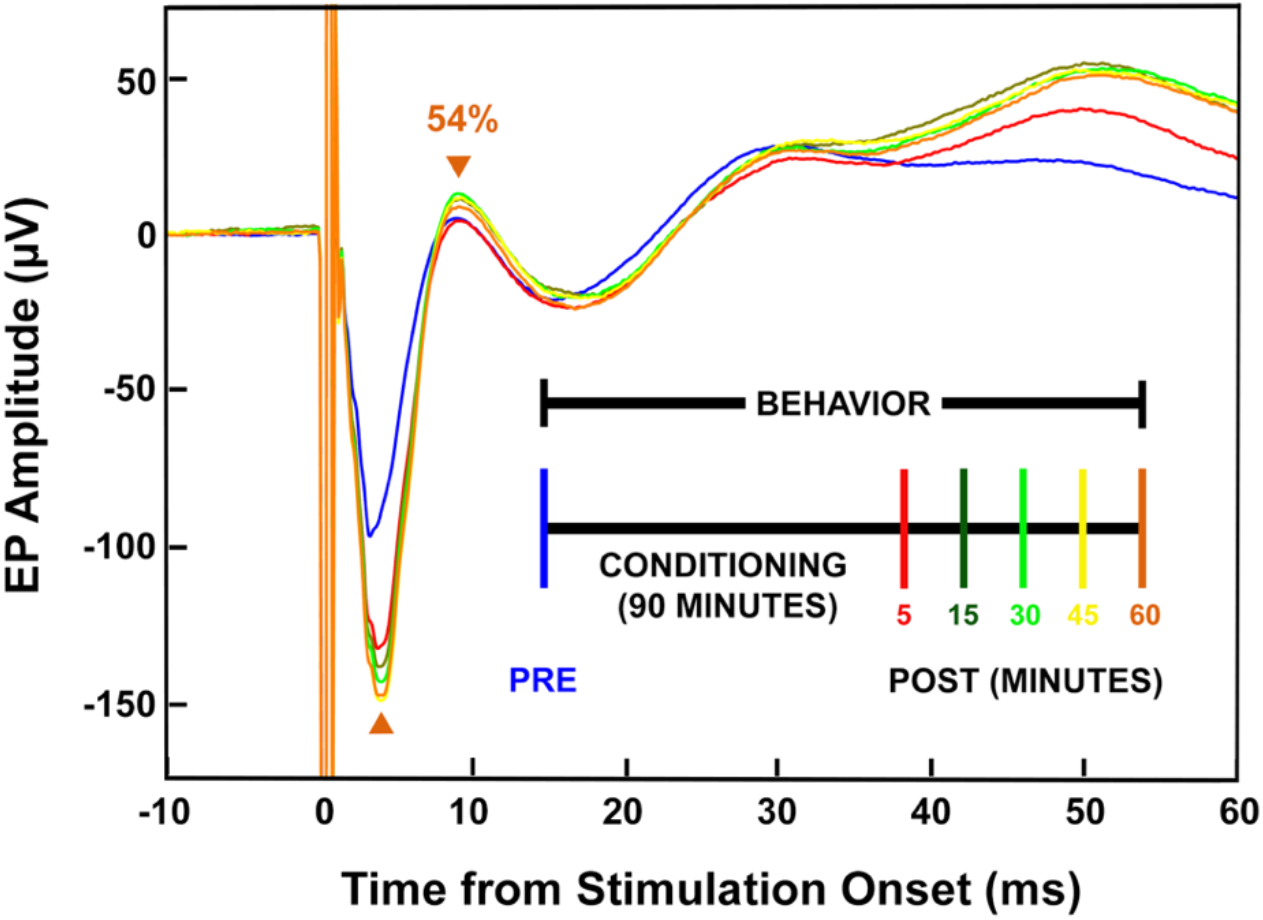
Postconditioning behavior modulates conditioning-induced cortical plasticity. StTAs of cortical responses were used to document changes in connectivity from Ntest to Nstim. Inset shows the time course of the experimental session. The EP amplitude was characterized as the size of the trough-to-peak (indicated by the orange arrowheads) deflection in the early component (≤ 15 ms) of the evoked response, which was likely the product of a monosynaptic connection from Ntest to Nstim (28). Color of the StTA curves in the plot corresponds to events in the inset timeline.

Delivery of movement-gated stimulation produced a 34% increase in the strength of the connection from Ntest to Nstim when assessed five minutes after the end of conditioning (*cf*. blue and red traces in Figure 2). Notably, the EP amplitude continued to grow in an incremental manner over time, while the monkey performed the wrist task, during the postconditioning period (instead of decaying back to the preconditioning level). At the 60-minute time point (orange trace), the EP was 54% larger compared to the preconditioning level (blue trace), suggesting a role for postconditioning behavior in modulation of conditioning-induced strengthening of cortical connections.

### Movement-gated conditioning combined with postconditioning behavior produces further strengthening of cortical connections

To assess the role of postconditioning behavior in modulation of conditioning-induced cortical plasticity and examine the time course of washout, we documented connectivity changes for two weeks post conditioning. During the two weeks, we conducted 2.5-hour behavioral sessions every weekday. Test stimuli were delivered at multiple time points during these sessions to assess the strength of the connection from Ntest to Nstim while monkeys performed the wrist target-tracking task. Note that no additional conditioning stimulation was delivered during these behavioral sessions.

Figure 3 shows the experimental timeline (Figure 3A) and connectivity changes (Figure 3C) at a pair of sites in the left primary motor cortex of monkey Y, shown in Figure 3B with Ntest in red and Nstim in green. Preconditioning motor outputs were assessed using trains of ICMS whose delivery at Ntest activated wrist extensors at a lower threshold intensity compared to flexor muscles, so extension-gated stimulation was delivered to Nstim during conditioning. Note that preconditioning delivery of ICMS trains at Nstim produced activation of both wrist flexors and extensors.

**Figure 3.**
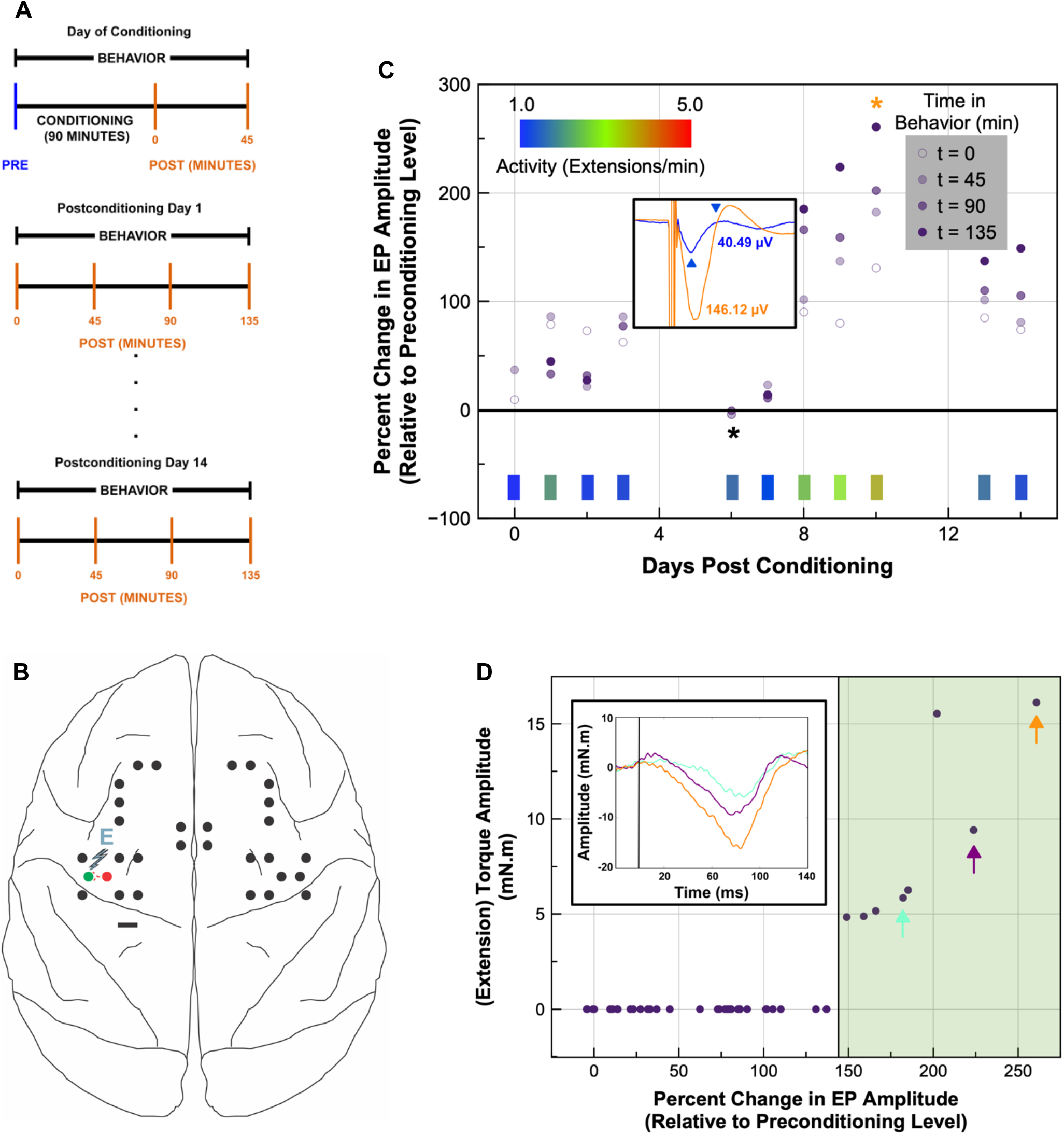
Movement-gated stimulation combined with behavior produces volitional strengthening of cortical connections. (A) Movement-gated conditioning was delivered on day 0 (for 90 minutes) and was preceded and followed by delivery of test stimuli on the same day (at two time points; 0, 45) and during behavioral sessions that occurred every weekday over the subsequent two weeks. Test stimuli were delivered at four time points (0, 45, 90, 135) during these behavioral sessions. (B) Schematic of the implant in monkey Y with the conditioning site pair, whose data are shown in (C), indicated in red and green. Extension-gated stimulation was delivered at Nstim (green site) during conditioning. E, extension; scale bar, 3 mm. (C) The percent change in EP amplitude was used to document the effect of combining conditioning and behavior on cortical connectivity during the 15-day experiment. Connectivity was assessed at the orange time points shown in the timeline in (A), represented here in different shades of violet. Color bars (at the bottom of the plot) overlaid on a graded three-color heatmap (shown in inset on top left) indicate the average frequency of wrist extensions during each behavioral session. The maximum and minimum values of the connectivity dataset are indicated by orange and black asterisks, respectively. Inset in the center of plot shows StTAs of the preconditioning EP (blue) and the EP evoked at the maximum gain (orange), with color-matched labels indicating the respective EP amplitudes. The EP amplitude was characterized as the size of the trough-to-peak (indicated by blue arrowheads) deflection, shown here for the preconditioning EP. The same trough and peak were used to characterize postconditioning amplitudes. Note behavioral sessions only occurred on weekdays; lack of data on days 4, 5, 11 and 12 is due to their occurrence over weekends. (D) Plot shows amplitude of stimulus-triggered extension torques as a function of cortical-connectivity changes. Inset shows StTAs of wrist torques (with flexion and extension responses represented by positive and negative deflections, respectively) at three selected cortical-gain values indicated in the main plot by color-matched arrows.

Successive performance of the wrist task for days after conditioning produced dramatic strengthening of the connection from Ntest to Nstim, as seen by an increase in the size of cortical potentials evoked at Nstim (Figure 3C). The postconditioning EP amplitudes in many cases were over 100% larger compared to the preconditioning level. Second, the larger postconditioning EPs coincided with higher frequency of wrist extensions, which, in this case, occurred on days 8, 9 and 10 post conditioning. During these three days, the size of the EP grew incrementally, both during each session and over the three sessions, from 90% (on day 8 at t = 0 time point; day 8/t = 0) to 261% (on day 10/t = 135; indicated by the orange asterisk), which was the maximum gain observed during the 15-day conditioning-and-behavior experiment. Substantial gains were also seen both before (on days 0, 1, 2 and 3) and after this three-day period. Third, there were changes in the size of the EP between the last (*i.e*., t = 135) time point of a behavioral session and the first (*i.e*., t = 0) time point of the subsequent session, possibly due to bidirectional homeostatic adjustments of synaptic strength (30, 31) during sleep (32, 33). This change was more pronounced when the latter session occurred after a weekend and resulted in a decrease in the strength of the connection towards preconditioning levels (*cf*. day 10/t = 135 and day 13/t = 0 in Figure 3C). Finally, the resultant cortical plasticity persisted for over two weeks with the size of the EP still 149% larger (or the connection 2.5-fold stronger) on day 14 (at t = 135 time point) post conditioning.

We also investigated the effect of combining conditioning and behavior on the late component of the Ntest-to-Nstim EP and the motor output of Ntest. At the maximum gain of 261% in the magnitude of the early component (see orange asterisk in Figure 3C), there was a concomitant increase of 315% in the amplitude of the late component of the EP (measured between 15 – 100 ms) relative to preconditioning level. Output effects were quantified with StTAs of flexion-extension wrist torques (16) evoked by delivery of the same test stimuli at Ntest that were used for documenting the strength of its connection to Nstim (in Figure 3C). Figure 3D shows the magnitude of torque responses evoked from Ntest as a function of postconditioning cortical-connectivity changes. Preconditioning test stimuli did not evoke any flexion or extension wrist torques (indicated by the origin of the plot), as expected, since currents that were subthreshold for movement were used for assessing connectivity and test stimuli were always delivered with the monkey’s wrist at rest. No postconditioning torques were evoked when cortical gains were less than 148%. Once this threshold was exceeded (marked by the green-shaded region in the plot), the same test stimuli evoked wrist torques. In concordance with delivery of extension-gated stimulation (Figure 3B), postconditioning test stimuli evoked extension (as opposed to flexion) torques whose amplitudes were strongly correlated to gains in cortico-cortical connectivity (Pearson’s correlation coefficient = 0.84 in the green-shaded region of the plot), demonstrating a reinforcement in the output of motor-cortical sites that was consistent with the potentiation of the synaptic connection between them.

### Conditioning alone or behavior alone are ineffective in modulating cortical connectivity

To delineate the individual contributions of conditioning and behavior, we performed two controls at the same pair of left motor-cortical sites (in monkey Y) shown in Figure 3B. The first control (performed three weeks after the end of the conditioning-and-behavior experiment) assessed the effect of behavior alone by documenting changes in Ntest-to-Nstim EPs for 15 days without delivery of any conditioning stimulation on day 0 (Figure 4A). The second control assessed the contribution of conditioning through delivery of extension-gated stimulation followed by documentation of connectivity changes without the monkey performing the wrist task after conditioning on day 0 and during subsequent days. Due to practical challenges associated with monkeys sitting in the experimental recording booth without doing a task or receiving any reinforcements, test stimuli were delivered only at the start of each postconditioning session (*i.e*., at t = 0; Figure 4C). As shown in Figures 4B and 4D, both behavior alone and conditioning alone produced small cortical-connectivity gains (≤ 35%). Additionally, there was a trend towards depression of the connection over time, especially in the second week, associated with both behavior alone and conditioning alone. These control experiments suggest that conditioning or behavior, in isolation, are relatively ineffective in modulating the strength of cortical connections.

**Figure 4.**
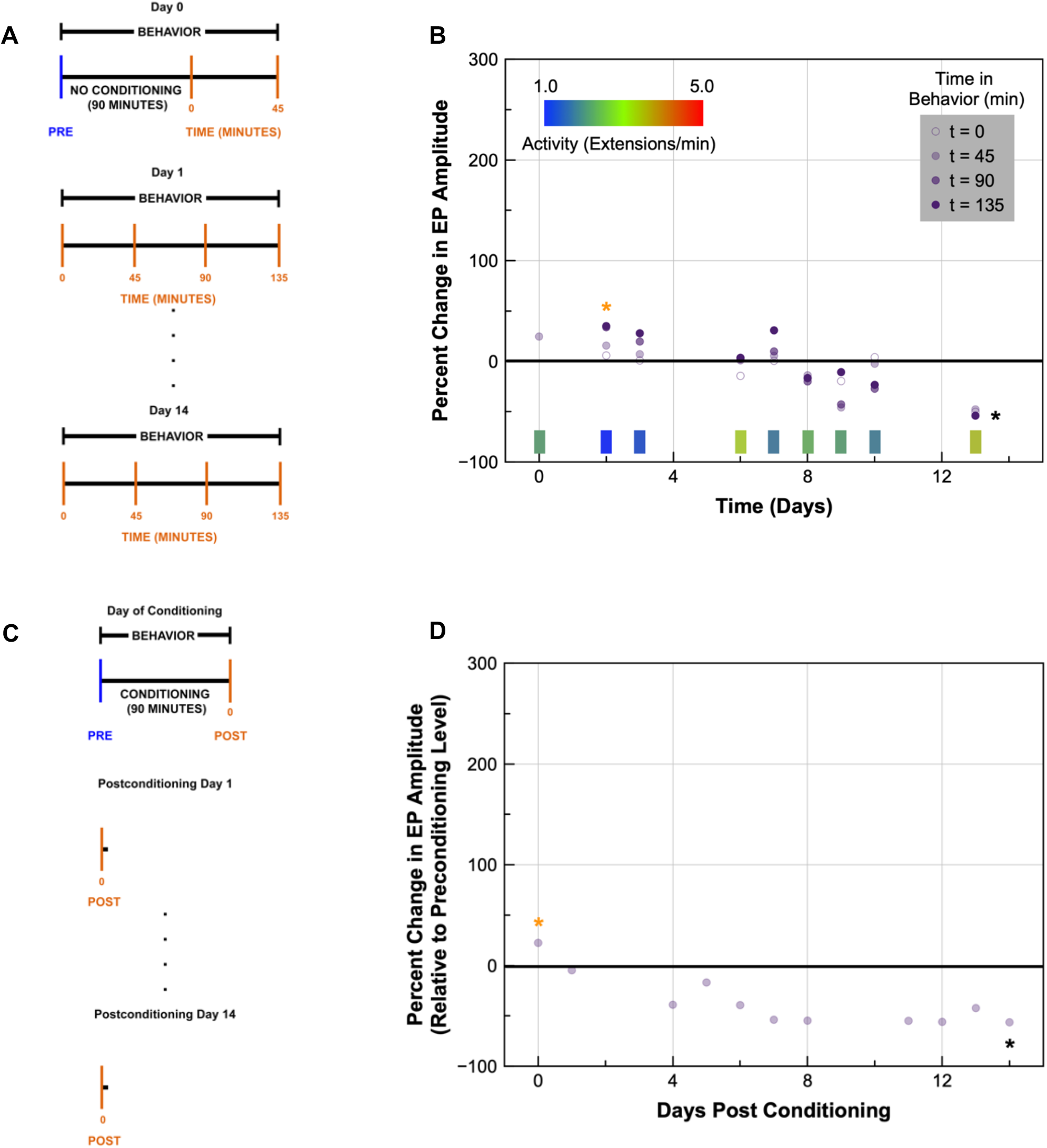
Behavior alone or conditioning alone are relatively ineffective in modulating cortical connectivity. (A) Timeline of behavior-alone control. Test stimuli were delivered on day 0 and during behavioral sessions (at four time points; 0, 45, 90, 135) that occurred every weekday over the subsequent two weeks. (B) The percent change in EP amplitude, relative to the initial assessment level denoted by ‘PRE’ in (A), was used to document connectivity changes with behavior alone. Activity-inset and color-bar descriptions are the same as Figure 3C. (C, D) Timeline and connectivity data for the conditioning-alone control. Data for both plots come from the same pair of left motor-cortical sites shown in Figure 3B. The maxima and minima of the two datasets in (B) and (D) are indicated by orange and black asterisks, respectively.

### Cortical strengthening produced by combining conditioning and behavior is direction specific

In a subset of experiments, we also explored connectivity changes in the reverse direction (*i.e*., from Nstim to Ntest). Here, test stimuli were delivered at Nstim to document changes in the strength of its connection to Ntest. We found that modulation of cortical EPs obtained by combining conditioning and behavior was direction specific, producing little to no changes in the reverse direction. Figure S1 shows connectivity changes in both the forward and reverse directions at a second pair of left motor-cortical sites during another 15-day conditioning-and-behavior experiment. Combining conditioning and behavior at this pair of sites (shown in Figure S1A) produced a maximum gain of 166% in the forward direction (on day 9/t = 0; indicated by the orange asterisk; Figure S1B) while the maximum gain in the reverse direction was 26% (which occurred on day 2/t = 135; also indicated by the orange asterisk; Figure S1C).

### Connectivity changes obtained with conditioning and behavior, delivered individually or together, across cortical sites

We investigated the effect of conditioning and behavior, delivered individually or together, on the strength of motor-cortical connections in the forward and reverse directions across eight site pairs in the two monkeys. Conditioning-and-behavior experiments were conducted at four site pairs in monkey Y and at three site pairs in monkey U (although only six out of those seven site pairs were tested in the reverse direction). The effect of behavior alone was tested at four site pairs in each monkey, while conditioning alone was tested at two site pairs in monkey Y and at three pairs in monkey U. Figures 5A and 5B document the connectivity changes across site pairs, showing all individual data points, whose means are shown in Figures 5C and 5D, respectively. Large cortical-connectivity gains were obtained only when conditioning was combined with behavior, and these gains were restricted to the forward direction (Figures 5A and 5B, green data points).

**Figure 5.**
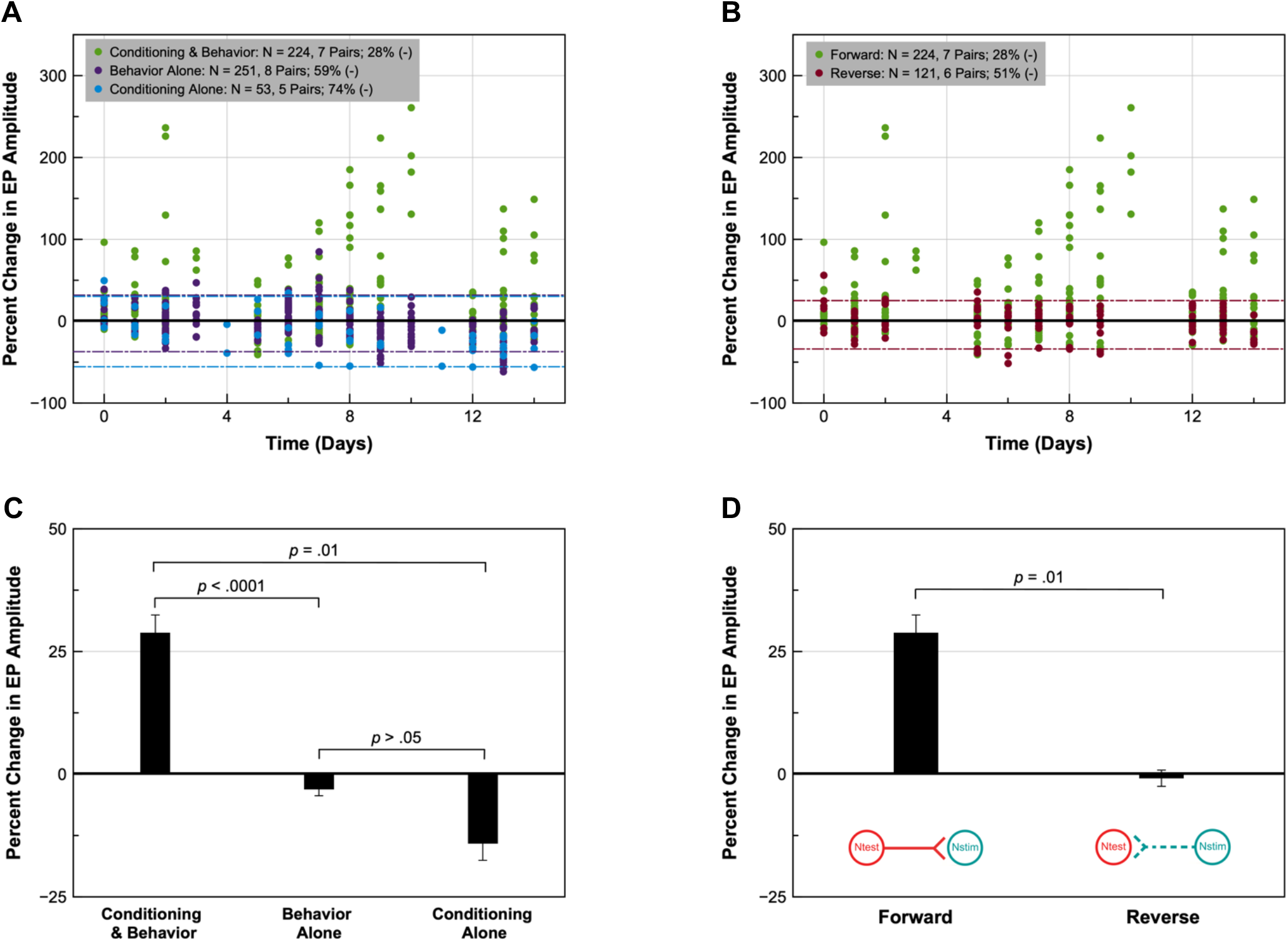
Connectivity changes obtained with conditioning and/or behavior across cortical sites. (A, C) The percent changes in EP amplitude in the forward direction produced by conditioning and behavior, delivered individually or together, are shown for all tested site pairs. Individual data points are shown in (A) with the corresponding means and significance levels shown in (C). Similarly, (B) and (D) compare connectivity changes observed with conditioning and behavior, delivered together, in the forward and reverse directions. Color-coded dashed lines show 5^th^ and 95^th^ percentiles for the corresponding control distributions in (A) and (B). Note that the 95^th^-percentile values for both controls are overlapping in (A). Sample sizes for all conditions are listed in (A) and (B), which includes both the total number of data points (denoted by ‘N’) and the corresponding number of cortical site pairs tested under each condition. (-) indicates percentage of changes with negative polarity (*i.e*., post < pre). Error bars in (C) and (D) represent standard error of the mean. *p*-values were obtained using linear-mixed-model analyses.

Due to repeated assessment of EP amplitudes over time at individual connections and presence of unbalanced data (arising from unequal sample sizes across groups), we implemented a linear mixed model to compare changes within and across intervention groups (34), details of which can be found in SI Appendix, Materials and Methods. The linear-mixed-model statistical analyses revealed a significant overall difference between the conditioning-and-behavior and behavior-alone groups (*F*_1,33.226_ = 20.462, *p* < .0001; effect size = 1.54, very large effect; Figure 5C) as well as a differential interaction of group type over time (*F*_49,273.248_ = 2.611, *p* < .0001), which was also confirmed individually in the two animals (monkey Y: *F*_30,123.364_ = 3.355, *p* < .0001; effect size = 1.76, very large effect and monkey U: *F*_30,115.119_ = 1.872, *p* < .01; effect size = 0.95, large effect). Changes obtained by combining conditioning and behavior were also significantly different from conditioning alone (*F*_1,17.220_ = 8.143, *p* = .01; effect size = 0.97, large effect; Figure 5C), while there was no significant difference between the behavior-alone and conditioning-alone groups (*F*_1,20.334_ = 4.141, *p* > .05; Figure 5C). Additionally, connectivity changes produced by conditioning and behavior, delivered together, in the forward direction were significantly larger than those in the reverse direction (*F*_1,27.984_ = 7.544, *p* = .01; effect size = 1.48, very large effect; Figure 5D). Details of effect-size calculation and thresholds used for interpretation of the magnitude of observed effects can be found in SI Appendix, Materials and Methods.

We also found that connectivity gains in the forward direction obtained immediately after conditioning (22.10 ± 1.38, mean ± standard error) were significantly smaller than the maximal changes (123.71 ± 2.62, mean ± standard error*; e.g*., see orange asterisks in Figures 3C, S1B, S2B and S2D) produced by further repetition of wrist movements, without delivery of any additional conditioning stimulation (*F*1,12.019 = 7.484, *p* = .02, linear mixed model using individual-trial data; effect size = 1.53, very large effect). A paired samples t-test further revealed that postconditioning StTAs immediately after conditioning (*i.e*., on day 0/t = 0) were not significantly different from preconditioning StTAs (*t*(6) = 1.826, *p* > .05). Conversely, postconditioning StTAs corresponding to the maximal changes, obtained through repetition of “conditioned” movements, were significantly larger than preconditioning StTAs (*t*(6) = 2.713, *p* = .03; effect size = 2.67, very large effect).

While conditioning and behavior, when combined, produced significant gains in the strength of forward connections, the magnitude and duration of cortical plasticity observed across site pairs were variable (*cf*. Figures 3C, S1B, S2B and S2D). Effect sizes, calculated at the maximal postconditioning changes obtained during the 15-day conditioning-and-behavior experiments, ranged from small to very large across connections and animals (monkey Y: 0.39 – 5.09 and monkey U: 0.64 – 2.75). There was a trend for inverse dependence of modifications on the initial strength of the cortical connection; larger potentiation occurred in weaker connections, while stronger connections were harder to modify. A similar observation was made by Poo and coworkers in their seminal paper on STDP (1). Furthermore, in some cases, gains were slowly accumulated over successive behavioral sessions (see Figures 3C, S1B and S2D), while they were accrued over faster timescales at other cortical site pairs. In a striking example, shown in Figure S2B, the strength of the forward connection increased, when conditioning was combined with behavior, from −5% to 237% within an individual (135-minute) behavioral session on day 2 post conditioning. However, the resultant plasticity was short-lived and extinguished over the weekend that followed the session (in contrast to the more persistent effect shown in Figure 3C).

Reproducibility of conditioning effects was assessed at a single site pair. We delivered both conditioning-and-behavior and conditioning alone twice at the same pair of left motor-cortical sites shown in Figure 3B. The interval between the two conditioning-and-behavior experiments was 4.82 months, while the two conditioning-alone experiments were performed 6.07 months apart. Qualitatively similar results were obtained across repetitions (*cf*. Figures 3C and S3A & Figures 4D and S3B).

Lastly, we assessed the directionality of all connectivity changes (relative to baseline levels) obtained with conditioning and/or behavior across the 15-day duration of our experiments (Figure 5A). We found that 28% of the changes obtained by combining conditioning and behavior were negative. In stark contrast, 59% and 74% of the changes were negative with behavior alone and conditioning alone, respectively, indicating higher levels of depression. A paired samples t-test further revealed that postconditioning StTAs corresponding to the minimal changes (*e.g*., see black asterisks in Figures 3C, S1B, S2B and S2D), obtained by combining conditioning and behavior, were not significantly different from preconditioning StTAs (*t*(6) = −1.811, *p* > .05). On the other hand, post-intervention StTAs corresponding to the minimal changes were significantly smaller than preintervention StTAs with both behavior alone (*Z* = −2.521, *p* = .01, Wilcoxon signed-rank test; effect size = 0.45, small effect) and conditioning alone (*t*(4) = −3.427, *p* = .03; effect size = 0.61, medium effect).

### Global effects of combining conditioning and behavior

STDP-based stimulation protocols have often been found to produce global modifications of synaptic strength, affecting connections beyond the site of induction (27, 35–39). To test site specificity of the cortical potentiation induced by combining conditioning and behavior, we documented changes in the strength of outgoing connections from Ntest and Nstim to other implanted sites. We found that the cortical gains observed with conditioning and behavior, delivered together, were not restricted to Ntest-to-Nstim, but the potentiation propagated outwards to affect all connections from the presynaptic site (Ntest; Figure 6A), while the strength of connections from Nstim to other implanted sites showed little change (Figure S4).

**Figure 6.**
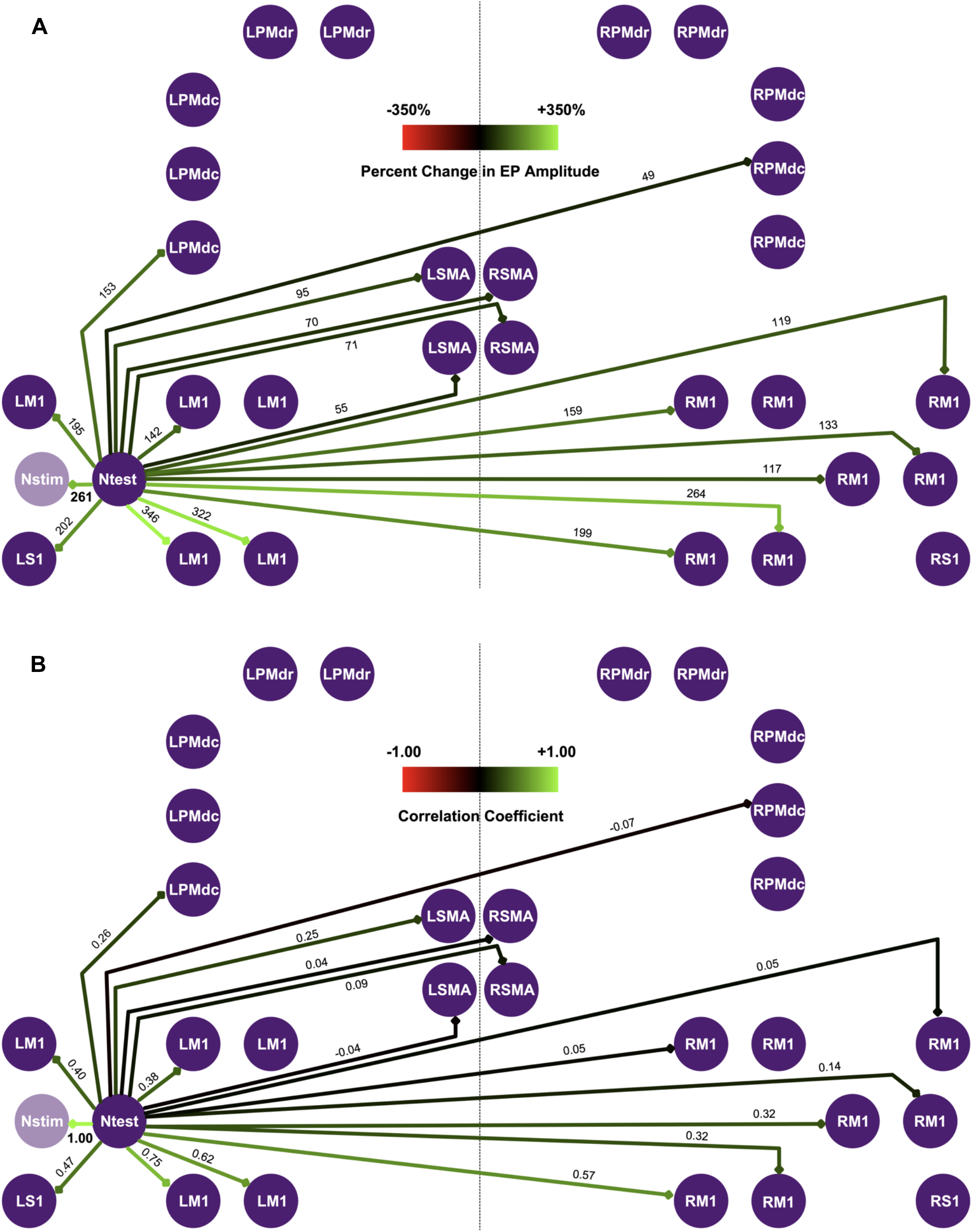
Global effects of combining conditioning and behavior. (A) The percent change in EP amplitude was used to document global connectivity changes produced by combining conditioning and behavior. Concurrent potentiation in the strength of all outgoing connections from the presynaptic site (Ntest) was observed when there was a 261% strengthening (indicated in bold; also see orange asterisk in Figure 3C) of the connection from Ntest to Nstim. Colored arrows denote the magnitude of connectivity changes imposed on a graded three-color (red-black-green) scale (inset on top); numbers indicate percent changes. (B) The same dataset in (A) was used to compute correlation coefficients between Ntest-to-Nstim EPs and EPs from Ntest to other sites. Colored arrows, here, denote the correlation coefficients imposed on the same graded three-color scale (inset on top), with numbers indicating coefficient values. For (A) and (B), the schematic represents the implant of monkey Y. Lack of arrows implies that there were no clear connections, as assessed through EPs in StTAs, between sites. M1, primary motor cortex; S1, primary somatosensory cortex; SMA, supplementary motor area; PMdr, rostral subdivision of the dorsal premotor cortex; PMdc, caudal subdivision of the dorsal premotor cortex. Prefixes L and R indicate the left and right hemispheres, respectively. Note that both Ntest and Nstim are sites in LM1.

In the example shown in Figure 6A, a 261% potentiation in the Ntest-to-Nstim connection (see orange asterisk in Figure 3C), produced by combining conditioning and behavior, resulted in a concurrent increase in the size of all outgoing EPs from Ntest. The magnitude of potentiation was variable, ranging from 49% to 346%, across sites, which spanned different cortical areas (primary motor cortex, primary somatosensory cortex, premotor cortex, supplementary motor areas) and both the left and right hemispheres. Importantly, gains were not correlated to the motor outputs of the cortical sites or their distance from Ntest or Nstim. Motor outputs across cortical sites ranged from flexion, extension, both flexion and extension to neither flexion nor extension. In a second example shown in Figure S5, a 237% increase in the strength of the connection from Ntest to Nstim (see orange asterisk in Figure S2B), similarly, produced concurrent potentiation in all outgoing connections from Ntest that ranged from 66% to 242%. Again, the sites spanned different cortical regions and both hemispheres, and potentiation was not related to their motor outputs or their distance from Ntest or Nstim. Second, potentiation of the Ntest-to-Nstim connection produced very small (bidirectional) changes in the strength of all outgoing connections from Nstim, including its connection to Ntest (Figure S4). Lastly, this pattern of global plasticity required induction of potentiation and was absent when there was no substantial change in the strength of the connection from Ntest to Nstim. As shown in Figure S6, a −1% change in the strength of Ntest-to-Nstim connection was accompanied by changes ranging from −38% to 41% across sites that received inputs from Ntest. Note all sites that received inputs from Ntest or Nstim, assessed through presence of EPs in StTAs, were included in these analyses.

To test whether the strengthening of outgoing connections from Ntest was due to an increase in the excitability of neurons, we computed the Pearson’s correlation coefficient between Ntest-to-Nstim EPs and EPs evoked from Ntest to a second site during individual trials (Figure 6B). Higher values of the correlation coefficient across sites would support a covariation in EP sizes due to a generalized increase in Ntest excitability, presumably produced by a greater response of neurons at Ntest to the test stimulation. In contrast, we found that the correlation coefficients were variable across sites, ranging from −0.07 to 0.75, suggesting that outgoing modifications in connectivity from Ntest were unlikely to have resulted from changes in excitability alone.

## Discussion

This study describes an activity-dependent stimulation protocol in which electrical stimuli gated by voluntary movements were used to produce simultaneous activation of neurons at motor-cortical sites in adult behaving monkeys. Movements corresponding to the motor output of a presynaptic site, during which neurons at that site were likely active, gated delivery of stimuli to a postsynaptic site in an effort to produce co-activation of cells at the two sites. Delivery of movement-gated stimulation resulted in small increases in the strength of cortical connections immediately after conditioning. Gains in cortical connectivity and motor output, induced by STDP-based conditioning approaches, have been reported before by our laboratory (6, 7, 16, 27) and other researchers (40–42). In these studies, conditioning-induced plasticity decayed over minutes to several days (depending on the duration and frequency of conditioning sessions). In contrast, when movement-gated stimulation was paired with further repetition of the movements that gated the conditioning stimuli, there were substantially larger increases in the strength of cortical connections, without any additional delivery of conditioning stimulation. Moreover, conditioning alone or behavior (used here to describe both voluntary movements and associated levels of behavioral modulators, such as attention and motivation) alone were relatively ineffective in modulating cortical connectivity. Second, the cortical plasticity produced by combining conditioning and behavior was directional, strengthening connections in the forward direction with little to no changes in the reverse direction. Lastly, there was a global, but selective, spread of plasticity from the conditioned sites that resembled a presynaptic pattern of propagation of potentiation, previously described in an *in vitro* study of STDP (35).

### Comparison with related activity-dependent stimulation protocols

The current conditioning paradigm is related to protocols used in our previous studies in which cortical or spinal stimulation was triggered by multiple motor unit action potentials (MUAPs) in forelimb-EMG recordings (16, 43). Lucas and Fetz showed that such MUAP-triggered cortical stimulation produced a shift in the ICMS-evoked movement representation of the presynaptic site (associated with the recorded triggering muscle) towards that of the postsynaptic stimulated site, presumably due to strengthening of the cortical connection from the pre- to the postsynaptic site (16). In neural-network simulations, the changes in intracortical connectivity were consistent with STDP mechanisms (44). In the second study, McPherson *et al*. improved forelimb-motor recovery in rats with incomplete cervical spinal-cord injury by delivering long-term MUAP-triggered spinal stimulation, which may have strengthened spared corticospinal connections to neurons below the lesion (43).

These two previous studies (16, 43) and the current investigation used signals related to muscle activity as a proxy for movement-related firing of motor-cortical neurons. However, it is worth considering the differences between MUAP-triggered and movement-gated stimulation with regards to the underlying plasticity mechanisms. MUAP-triggered stimulation is designed to synchronize stimulus-driven action potentials of postsynaptic neurons with spikes of presynaptic cortical cells that are co-activated with EMG in the triggering muscle. This coupling is tightest between corticomotoneuronal (CM) cells and motoneurons of their target muscles in primates (45), due to their monosynaptic connections. CM-cell spikes are also synchronized with firing of other cortical neurons, as shown by peaks in their cross-correlograms (46). Spike-triggered averages of EMG demonstrate that CM cells fire 6 - 25 ms before MUAPs of their target muscles (45), indicating that MUAP-triggered cortical stimulation is within the window for STDP (1) to strengthen intracortical synapses of CM and synchronized cells.

Despite evidence for temporal coupling (45), the directional tuning of CM cells and their target motor units is not always the same (47, 48). Moreover, many populations of last-order premotor neurons contribute to motor-unit firing in addition to CM cells (49), making the timing between CM-cell and motor-unit firing probabilistic. The relative timing of firing of non-CM motor-cortical cells, synchronized with CM cells, and motor units is expected to be broader, given the intervening synapses (making the coupling weaker). When using trains of ICMS to document cortical representation, as in the Lucas and Fetz study (16), it is also important to take into account that repetitive stimulation activates additional pathways through temporal summation, making the extent to which CM-cell activation evokes the output effects unknown and quite possibly minor. Despite these complexities, results in the Lucas and Fetz study are consistent with an STDP mechanism (16, 44), suggesting that the population of cortical neurons that produced the shift in the direction of ICMS effects had a statistically higher probability of firing less than ~30 ms before the MUAP-triggered stimulus pulses that activated their postsynaptic targets in the motor cortex.

In contrast, it is difficult to attribute the plasticity produced by movement-gated stimulation to a similar STDP mechanism. Although the firing rates of wrist-muscle MUAPs and wrist-related neurons in the primary motor cortex are simultaneously elevated during movement, the relative timing of conditioning-stimulus pulses and presynaptic cortical spikes would be essentially random because movement-gated stimuli were delivered in a train at a constant frequency. Thus, the number of presynaptic cortical spikes occurring before and after individual conditioning-stimulus pulses in the current study were likely equal, leading to as many weakening as strengthening STDP events. It is possible that the effects of the conditioning stimulation itself during the 1.6-second train, interacting with the complex excitatory-inhibitory circuits in the motor cortex, establish spike-stimulus timings that promote STDP (50). However, since the population of cortical cells contributing to the measured variable (*i.e*., EP amplitude) is unknown (*e.g*., the proportion of cells with CM *vs*. only cortico-cortical projections), it is difficult to formulate a specific hypothesis about potential circuit mechanisms. Alternatively, primarily non-Hebbian forms of plasticity may underlie the changes in connectivity reported in the current study. At the very least, increases in connection strength that occur long after delivery of conditioning stimulation must involve mechanisms other than STDP (which are discussed below). Neuromodulatory systems, for example, can influence STDP rules—by acting *via* acetylcholine, monoamines and other signaling molecules—and bridge the gap between the timing of spikes and their behavioral outcome (51, 52).

Lastly, the output measures used to assess the effect of conditioning were different across the three protocols, namely amplitude of cortico-cortical EPs (used here), ICMS-evoked torques (16) and motor performance (43), making it difficult to infer the relationships between the underlying plasticity mechanisms. In addition, the frequency and duration of the conditioning sessions and the timeline for testing the effects of conditioning were variable across the three studies. The largest changes in connectivity in the present study occurred with further performance of a motor task after a brief period of conditioning. Similar (and longer) conditioning periods were used by Lucas and Fetz (16), but the effect of postconditioning behavior on ICMS-evoked torques was not evaluated. Rats in the McPherson *et al*. study received many hours of daily MUAP-triggered stimulation for months (43); this led to sustained postconditioning changes, which could involve mechanisms similar to those in the current study.

### Role of behavior in cortical plasticity

Our results show that movement-dependent stimulation creates a plastic landscape in which repetition of the behavioral context presented during conditioning drives cortical strengthening long after conditioning has ended. In contrast, conditioning alone was relatively ineffective in modulating the strength of cortical connections. Moreover, the size of the cortical gains produced by combining conditioning and behavior was often positively correlated with the monkey’s wrist-activity levels (*e.g*., Figure 3C, days 8, 9, 10), suggesting a role for motivated behavior in cortical plasticity. In a relevant study supporting this argument, Nishimura and coworkers used Granger-causality analysis to demonstrate signal flow in the high-gamma frequencies from the nucleus accumbens to the primary motor cortex in the first 4 – 5 weeks following a cervical spinal-cord injury in macaque monkeys (18). This interaction was critical to rehabilitation-mediated recovery of forelimb-motor function. Pharmacological inactivation of the nucleus accumbens, involved in the regulation of motivation-driven effort (53–55), during this early phase post injury abolished the high-gamma activity, leading to deficits in recovery of finger dexterity. This study provides strong support for the role of motivated behavior in cortical plasticity and functional motor recovery.

Two other studies provide critical support for the role of behavior in synaptic plasticity. Ahissar *et al*. found that strengthening of cortical connections, produced by a conditioning protocol that co-activated neurons involved in those connections, was strongly dependent on the behavioral context of the stimuli that induced such modifications (12). Correlated activity of neurons, while necessary, was not sufficient for the induction of plasticity. Second, Perez and coworkers found that volitional activity during a paired stimulation protocol enhanced corticospinal transmission in humans with spinal-cord injury (19). Transmission was significantly improved when the paired stimulation was combined with relevant isometric movements compared to delivery of stimulation at rest, underscoring the role of volitional activity, which occurs at a very different timescale from STDP (8), in boosting synaptic plasticity.

Synaptic tagging (56–58) and changes in neuronal excitability (59) are two mechanisms by which synapses and neurons “primed” during the plasticity-induction state can be marked for further modifications. Such mechanisms may be involved in the effects observed with conditioning and behavior, delivered together, in the present investigation. In this scenario, synapses between neurons at Ntest and Nstim that are active during wrist flexion or extension may have been tagged by movement-gated stimulation and further strengthened by repetition of those movements in a use-dependent manner. In the absence of conditioning stimulation, no such priming occurs and performance of an already-learned motor task does not produce an appreciable increase in the strength of cortical connections associated with the task, indicating that cortical plasticity, while functionally relevant, is unnecessary for continued task performance (60). Similarly, tagging synapses, through movement-gated stimulation, followed by nonuse conditions synapses whose strength quickly decays to baseline levels.

### Interplay between various forms of plasticity

Our study shares features of STDP, such as directional modulation of synaptic strength (1–5) and a presynaptic spread of potentiation (35). However, connectivity changes observed in our experiments were likely also affected by non-Hebbian forms of plasticity, such as homeostatic plasticity (driven by cell-wide, and not synapse-specific, mechanisms such as global synaptic scaling; 61, 62) and slow-onset synaptic potentiation (56, 63), and the interactions between Hebbian and non-Hebbian forms of plasticity (30, 31). For example, there were bidirectional changes in the strength of cortical connections, produced by combining conditioning and behavior, between the last time point of a behavioral session and the first time point of the subsequent session (*e.g*., Figure 3C), possibly due to homeostatic adjustments of synaptic strength that occurred during sleep (32, 33). The changes were more pronounced when the latter session occurred after a weekend, and it resulted in a general decrease in connection strength towards preconditioning levels. Sleep is known to produce bidirectional changes in connection strength (32, 33). Induction of plasticity during awake states is often accompanied by secondary modifications during sleep through various forms of metaplasticity (64, 65), which refers to neuronal changes that influence the capacity for subsequent synaptic plasticity. Synaptic tagging (56–58) and changes in neuronal excitability (59) are, once again, implicated as mechanisms by which neurons “primed” during awake states can be marked for further modifications during sleep, thus providing a bridge between plastic changes across brain states. Second, this interaction between different forms of plasticity may also be reflected in the bidirectionality of connectivity changes (relative to baseline levels) produced by conditioning and behavior, delivered individually or together, over the 15-day course of the experiments (Figure 5A). The gains produced by combining conditioning and behavior likely offset the percentage of negative changes that were observed in the behavior-alone and conditioning-alone controls. Third, this interplay between gains produced by conditioning-and-behavior and homeostatic mechanisms, seeking to compensate for increases induced by activity-dependent stimulation in uninjured animals, may have also resulted in the lack of plateau in cortical-connectivity gains that was observed in our study. If synapses are potentiated and the resultant increases in synaptic strength are very long-lasting, more resources (neurotransmitters, receptors, vesicles, second messengers) would be needed to maintain such high levels of potentiation, which could potentially overwhelm the metabolic capacity of the system to stably sustain itself (61). These gains may, however, be more persistent in an injured environment where neuronal pathways are weakened and interventions seek to drive connection strength back to baseline levels.

### Propagation of cortical plasticity

The global pattern of plasticity seen in our study has been observed before by Poo and coworkers who found presynaptic propagation of longterm potentiation (LTP), produced by correlated pre- and postsynaptic activation, from the synapse of induction (35). In their study, which was conducted in sparse hippocampal cultures consisting of 3 – 4 neurons, LTP propagated retrogradely to glutamatergic synapses on the dendrites of the presynaptic neuron and laterally to those made by its axonal collaterals onto other glutamatergic cells. No lateral or forward propagation of LTP to or from the postsynaptic neuron or secondary propagation to synapses not directly associated with the pre- or postsynaptic neurons was observed. Also, as observed in our study (Figure S6), propagated potentiation required the induction of LTP. However, we have only investigated plasticity propagation in outgoing connections from the pre- and postsynaptic sites (Figures 6A, S4 and S5). Experimental time constraints associated with delivering sequential (test) stimulation at multiple sites precluded measurement of changes in incoming connections to the pre- and postsynaptic sites or secondary connections. Nonetheless, results from our experiments, conducted in behaving monkeys, bear remarkable similarities to some of the features of the plasticity-propagation pattern observed in the *in vitro* STDP study (35). Such selective propagation was found to be mediated by a retrograde messenger that rapidly crosses the synapse where potentiation was induced, signaling presynaptic modifications that produce a global, yet specific, pattern of connectivity changes at a timescale similar to plasticity induction (66, 67).

### Conclusions

Our results demonstrate that movement-dependent conditioning combined with repetition of the gating movements can produce significant strengthening of connections in the motor cortex of adult behaving monkeys. They indicate a crucial role for behavior, specifically voluntary movements and motivation, in modulating Hebbian-like plasticity. Physical rehabilitation and use-dependent movement interventions, such as body-weight-supported treadmill training (68) and constraint-induced therapy (69), have been widely explored as treatment options after neurological injuries, but they typically, and at best, lead to partial recovery of motor function. Our data provide strong support for combining movement-gated stimulation with such use-dependent physical therapies for enhancing motor recovery after a stroke or spinal-cord injury. Since our stimulation protocol uses a non-invasive gating signal (*i.e*., movement), subsequent studies will assess its clinical applicability by using less-invasive surface electrodes or superficial scalp electrodes for delivery of conditioning stimulation. Augmenting motivation through electrical stimulation of dopaminergic neurons in the nucleus accumbens or the midbrain may further enhance motor recovery promoted by movement-gated stimulation and physical rehabilitation (70), and exploration of such combinatorial interventions may represent important future directions. They will also help dissect the role of behavior, and behavioral modulators, in synaptic plasticity.

## Materials and Methods

All animal-handling, training and surgical procedures were approved by the University of Washington’s Institutional Animal Care and Use Committee, and they conformed to the National Institutes of Health’s Guide for the Care and Use of Laboratory Animals. For detailed descriptions of (1) animals, (2) behavioral training, (3) fabrication of cortical implants and surgeries, (4) EMG electrodes, (5) recordings and stimulations, (6) documentation of cortico-cortical connectivity and motor outputs, (7) delivery of conditioning and/or behavior, (8) connectivity analyses, and (9) statistics, refer to the *SI Appendix*.

## Acknowledgments

We thank Larry Shupe and Jatin Sonavane for providing programming, hardware and software assistance. We also thank Rebekah Schaefer, Andrew Bogaard and Robert Robinson for assistance with animal care, handling, training and surgeries. This work was supported by the National Institutes of Health (RR00166 and NS12542) and the National Science Foundation Center for Neurotechnology (EEC-1028725).

## Supporting Information

### Materials and Methods

All animal-handling, training and surgical procedures were approved by the University of Washington’s Institutional Animal Care and Use Committee, and they conformed to the National Institutes of Health’s Guide for the Care and Use of Laboratory Animals.

#### Subjects

Experiments were performed with two adult male *Macaca nemestrina* monkeys (1): Y (6.6 – 7.3 years old; 11.2 – 12.8 kg) and U (8.6 – 9.2 years old; 12.3 – 13.2 kg).

#### Behavioral training

Monkeys were trained to sit in a primate chair in a sound-attenuating recording booth, in front of a computer monitor, and perform a one-dimensional center-out force target-tracking task in which isometric wrist torque controlled the position of a cursor on the screen. The center target represented the “zero force” or neutral position of the wrist. Peripheral target positions alternated between flexion and extension and were presented in a random order with equal probability. Target placement on the screen determined the required direction and magnitude of wrist torque. Using their right hands, which were restrained in a manipulandum, monkeys were required to move towards and hold the cursor inside presented targets for 1.6 seconds (by exerting isometric force). Monkeys were rewarded with fruit sauce at a variable 1:1.5 reinforcement ratio after successful completion of trials. The manipulandum contained force transducers that measured wrist torques, which were recorded during experimental sessions. Additionally, detailed wrist-activity logs were saved at the end of each session. Training was complete when monkeys learned to hold the cursor inside presented targets for the duration of the hold time and performed the motor task consistently for approximately three hours, which was the typical duration of our experiments.

#### Cortical implant and surgery

After learning the motor task, monkeys received chronic bilateral (symmetrical) implants, consisting of custom-made arrays of dual electrodes (whose design is described in 2) with paired intracortical and surface Pt/Ir microwires. Briefly, bare (or uncoated) 127-μm diameter Pt/Ir microwires (A-M Systems, 767700), cut to two different lengths (3 mm for the surface electrodes and 5 mm for the intracortical electrodes), were soldered to 32-gauge insulated lead wires. Next, the soldered wires were coated with a 5 – 10 μm thick film of parylene C using chemical vapor deposition (3), at the Washington Nanofabrication Facility, which served to insulate both the Pt/Ir microwires and the solder joints. The parylene-coated wire assemblies were then deinsulated (using a scalpel or current-blasted) to expose the Pt/Ir tips until electrode impedances were less than 150 kΩ. This was followed by pairing of the intracortical and surface electrodes using polyimide tubing. The tips of the 3-mm and 5-mm Pt/Ir microwires were placed ~0.5 mm and 2.0 – 2.5 mm from the edge of the polyimide tube, respectively. Flowable silicone (Dow, 734) was squirted into the tube to hold the paired electrodes together. Lastly, the back ends of the lead wires were soldered to electronic connectors (Digi-Key Electronics) to complete the implant. The exposed length of the intracortical microwires was chosen to target sites in layer V of the sensorimotor cortex (which contains the cell bodies of pyramidal neurons; 4), while the shorter microwires rested on the surface of the brain.

All surgeries were performed under isoflurane anesthesia and aseptic conditions. During the cortical-implant surgery, an incision was made along the midline of the scalp, and muscle and connective tissue were resected to expose the skull. The dual electrodes were advanced individually through 1.3-mm diameter burr holes drilled in the skull with a stereotaxically-positioned drill. 15 sites in the sensorimotor cortex, involved in the planning and execution of wrist and hand movements (5), were implanted per hemisphere with dual electrodes in each monkey. Electrodes were distributed around the central sulcus in monkey U to target sites in the primary motor and somatosensory cortices (Figure S2C). In addition to targeting these two cortical areas, monkey Y received dual electrodes in the premotor cortex and supplementary motor areas (Figure 3B). Adjacent implanted sites were at least 2 mm apart. After advancing the dual electrodes, the burr holes were sealed with a thin layer of dental acrylic. The two connectors (one per hemisphere) were then cemented on the skull, also using dental acrylic. Next, titanium skull screws were placed remotely from the implanted areas to serve as grounds and references. All electrode lead wires as well as the ground and reference screws and wires were covered using acrylic. Lastly, a custom-made 2.5-inch diameter titanium head chamber, stabilized using additional titanium screws and held in place with acrylic, was used to protect the skull-mounted connectors. Animals received analgesics and antibiotics postoperatively.

#### EMG electrodes

Pre-gelled disposable surface patches with silver/silver chloride electrodes, placed on either side of the right forearm, were used to record EMG from wrist flexors and extensors. A sleeve reinforced the adhesion of the patches to freshly-shaved skin and held them in place during recordings.

#### Recordings and stimulation

Skull-mounted connectors were interfaced with two 32-channel headstages of a Cerebus™ data-acquisition system (Blackrock Microsystems), which was used for recording cortical LFPs from both hemispheres and evoked wrist torques. A third headstage was used for recording EMGs (also with the Cerebus™ system). Signals from all cortical electrodes were recorded single ended (relative to skull-screw grounds). Cortical signals were sampled at 10 kHz and 16-bit resolution using a low-pass 2.5-kHz filter. EMG signals were recorded differentially (at the same settings) with one electrode referenced to another on the same muscle. Wrist torques were sampled at the same frequency (unfiltered). Data from the amplifiers and analog inputs were streamed to a computer and visualized in real time using the Raster and Oscilloscope features in the Central graphic user interface provided by Blackrock. Bipolar cortical stimulation (with the intracortical electrode referenced to the paired surface wire) was delivered using a constant-current Nuclear Chicago stimulator. 200-μs wide biphasic symmetrical pulses (with zero delay between them) were used during stimulation. A custom-built relay device, fabricated at the University of Washington Instrumentation Services Core, interfaced with both the headstages and the stimulator and was used to switch remotely between recording and stimulation while monkeys sat in the recording booth. A trigger signal, routed from the stimulator into one of the analog inputs of Blackrock, was sampled simultaneously and was used to align recordings to generate StTAs of evoked cortical, muscle and torque responses.

#### Documentation of cortico-cortical connectivity and motor outputs

To identify connectivity between cortical sites, single-stimulus pulses were delivered at one site, through dual electrodes, and stimulus-evoked responses (or EPs) were recorded at all other implanted cortical sites (across both hemispheres). Stimuli were biphasic with the negative phase leading on the intracortical wire and the positive phase leading on the surface wire of the dual electrode. Stimuli used for assessing cortical connectivity were delivered at 4 Hz in a series of increasing current amplitudes, typically ranging from 0 – 300 μA, to identify the “EP threshold,” which was defined as the minimum current amplitude that evoked a clear EP (see Connectivity analyses below for more details).

Wrist-motor outputs were assessed through EMG responses evoked using 300-Hz trains of 3 – 7 (200-μs wide) biphasic ICMS pulses. Current amplitudes ranged from 50 – 300 μA and were increased in 50-μA increments. Stimulus-evoked responses in the flexor and extensor muscles, so called motor-evoked potentials (MEPs), were used to characterize the motor outputs of cortical sites. We determined the lowest current intensities at which MEPs appeared. Higher intensities often activated both flexors and extensors. Motor outputs were also characterized by measuring the amplitude of stimulus-evoked flexion and extension wrist torques, which were recorded simultaneously with EMG and cortical signals.

Stimuli used for assessing cortical connectivity or motor outputs, termed as test stimuli, were always delivered with the monkey’s wrist at rest. This was achieved by gating stimuli by cursor hold inside the center target during the center-out wrist task, which imposed a duty cycle with an interval equal to the travel time from and to the center target plus the hold time inside the peripheral (flexion/extension) target.

#### Delivery of conditioning and/or behavior

Reciprocal connections of two cortical sites (Ntest and Nstim) were identified *via* EPs (Figure 1B). Movement-related activity at the cortical sites was inferred from the motor outputs elicited by stimulation of those sites (Figure 1A). Conditioning stimulation was delivered to Nstim during movements that presumably activated neurons at Ntest to produce co-activation of the two sites. To achieve this, delivery of conditioning stimuli to Nstim was gated by cursor hold inside the flexion or extension target (depending on the motor output of Ntest). For example, if the motor output of Ntest was flexion, the monkey received flexion-gated stimulation at Nstim while the cursor was held inside the flexion target, as shown in Figure 1C. Movement-gated stimulation was delivered at 10 Hz, at an intensity that was subthreshold for movement, for a total of 90 minutes. In the above example, if the monkey held the cursor for the entire hold time (of 1.6 seconds), 16 (bipolar and biphasic) conditioning stimuli would be delivered per wrist flexion. A total of 8142 ± 688 (mean ± standard deviation) stimuli were delivered across sites during 90 minutes of conditioning. Delivery of conditioning was (preceded and) followed by delivery of test stimuli (at 4 Hz; Figure 1D), which were also subthreshold for movement, at both Ntest and Nstim to document changes in motor output and cortical connectivity (in both the forward and reverse directions).

Figure 3A shows the timeline of conditioning-and-behavior experiments. Conditioning was delivered on day 0 (for 90 minutes) and was preceded and followed (at two time points; 0, 45 minutes) by delivery of test stimuli on the same day and during behavioral sessions that occurred every weekday over the subsequent two weeks. Test stimuli were delivered at 3 – 4 time points (0, 67.5, 135 or 0, 45, 90, 135 minutes) during these behavioral sessions to test the strength of forward connections. Reverse connections were only tested twice (0, 135 minutes) during behavioral sessions to reduce the total amount of stimulation at Nstim (which also received the conditioning stimuli). Monkeys performed the flexion-extension target-tracking wrist task on the day of conditioning and during the following behavioral sessions. The frequency of movements that gated the conditioning stimuli was doubled during the conditioning and behavioral sessions to encourage use of the conditioned pathways. To assess the contribution of behavior alone on the observed plasticity, control experiments involved testing the strength of cortical connections at the same time points without delivery of any conditioning stimulation. Monkeys performed the target-tracking wrist task on all days (Figure 4A). Frequency of flexion or extension movements was doubled during these sessions to serve as identical controls for the corresponding conditioning-and-behavior experiments. A second control involved delivery of conditioning stimulation followed by connectivity testing without the monkey doing the wrist task post conditioning and during the following sessions. Due to practical challenges associated with monkeys sitting in the recording booth without doing a task or receiving any reinforcements, test stimuli were only delivered at the start of each session (*i.e*., at the t = 0 time points; Figure 4C) for this control.

Multiple Ntest sites were often included in the same experiment as long as they had reciprocal connections with the site selected as Nstim. To standardize the proportion of test and conditioning currents delivered across sites, we determined EP thresholds in both directions. Currents used for testing were ~2.0 x forward or reverse EP threshold for the forward and reverse connections, respectively, whereas ~3.5 x reverse EP threshold was used for conditioning.

#### Connectivity analyses

EP amplitudes were used for documenting the strength of cortical connections. Individual trials were aligned on stimulus onset (using the trigger signal from the stimulator) and grouped together. All trials were inspected and those with movement artifacts were removed. From the remaining trials, StTAs were generated from 20 ms before to 100 ms after stimulus onset. To be classified as an EP, the trough-to-peak (or peak-to-trough) deflection post stimulus (in the StTA) had to exceed two standard deviations of the baseline signal (calculated as the mean of samples between 5 – 20 ms preceding stimulus onset); otherwise, it was assumed that there was no evoked response at the recording site. Second, this response had to occur within 2.5 – 15.0 ms post stimulus. Lastly, we also looked for a clear separation from the stimulus artifact (which typically returned to baseline levels within 1.5 ms). The EP amplitude was then quantified by subtracting the largest trough from the largest peak in this early component (2.5 – 15.0 ms) of the mean evoked response (*e.g*., see Figure 2, arrowheads). Cortical-connectivity changes were quantified as percent changes in the average EP amplitude after conditioning and/or behavior relative to pre-intervention levels. Latency of the EP features in StTAs could vary slightly, so care was taken to ensure that the same peak and trough were used for a given EP across time points. Same rules were applied for assessing changes in the late component of the evoked response, except the latency range was 15 – 100 ms.

Changes in motor output were assessed by documenting the amplitude of stimulus-evoked flexion and extension wrist torques (in StTAs). Individual trials were, once again, aligned on stimulus onset and averaged to generate StTAs of wrist torques (Figure 3D). Torque responses were monophasic and were wired such that flexion and extension responses were represented by positive and negative deflections, respectively. The peak/trough latencies of evoked torques ranged between 71.7 – 87.0 ms in our experiments. Changes in motor output were also characterized by documenting the size of flexor and extensor MEPs in forelimb-EMG recordings.

In addition to characterizing changes in cortical connectivity at the conditioned site pairs, we documented changes in the strength of all outgoing connections from Ntest and Nstim in conditioning-and-behavior experiments. Second, we evaluated the covariation between Ntest-to-Nstim EPs and EPs evoked from Ntest to a second site during individual trials using Pearson’s correlation coefficient (Figure 6B).

All StTA analyses were performed using custom code written in MATLAB (MathWorks, Inc.).

#### Statistics

The order of interventions (conditioning and/or behavior) was randomized across sites. When behavior/conditioning alone followed conditioning-and-behavior at the same cortical site pair, we waited for a minimum of three weeks between the end of conditioning-and-behavior and the start of the subsequent control to allow for washout of gains. All changes in EP amplitudes were expressed relative to preconditioning levels, except for the behavior-alone control where the changes were relative to the initial assessment of the EP amplitude. Changes are reported as mean ± standard error. We also calculated the 5^th^ and 95^th^ percentiles of control distributions (which are shown in Figures 5A and 5B).

Due to the presence of correlations in EP amplitudes associated with repeated measurements at a given cortical connection and presence of unbalanced data, we implemented a linear mixed-effects model (6), in IBM SPSS Statistics (version 27), of changes in EP amplitude (which was our dependent variable) as a function of group, time, and the interaction between group and time (as our fixed effects). Cortical connections were included as a random effect. Compound symmetry or a first-order autoregressive (with homogeneous variances) repeated covariance structure was used, depending on which model fit the data better. Model fit was assessed using Hurvich and Tsai’s Akaike and Schwarz’s Bayesian information criteria, with lower values indicating better fit. Type III sum of squares (with intercepts included) was used to model the fixed effects. Satterthwaite’s method was used to estimate the degrees of freedom and generate *p*-values for the mixed models. Significance for all statistics was set at the .05 level.

For comparisons that involved the conditioning-alone group (in which EP amplitudes were assessed only at the start of each postconditioning session) with conditioning-and-behavior or behavior alone, we compared the t = 0 changes obtained with conditioning alone to the EP changes at the last time point of every behavioral session to ensure that a similar number of data points were included when comparing different intervention groups. The last time point was chosen to account for the effect of behavior in the conditioning-and-behavior and behavior-alone experiments. Similarly, when comparing changes obtained by combining conditioning and behavior in the forward *vs*. reverse directions, we used the first and last time points of every behavioral session since reverse connections were only assessed at these two time points.

Paired samples t-test was used for pre-post comparisons. Before running paired t-tests, we verified that the pre-to-post differences followed a normal distribution. Normality was assessed using the Shapiro-Wilk test. Wilcoxon signed-rank test, which is the nonparametric equivalent of the paired t-test, was used when the data were not normally distributed. All statistical analyses were performed in IBM SPSS Statistics (version 27).

In addition to assessing statistical significance, we also quantified the magnitude of the effect using the ‘effect size’ metric, which is the standardized difference of the mean of two groups, according to the following equation (7).

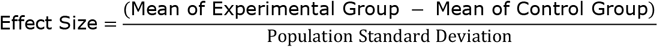

Population standard deviation was estimated by pooling the (sample-size-weighted) standard deviations of the experimental and control groups, using the equation below, when standard deviations of the two groups were similar. Otherwise, the standard deviation of the control group was used. These metrics correspond to Hedges’ g (8) and Glass’ delta (9) effect-size indices, respectively.

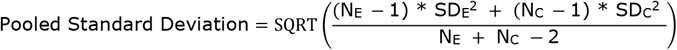

Here, N_E_, SD_E_, N_C_ and SD_C_ are the sample sizes and standard deviations of the experimental and control groups, respectively. Lastly, effect-size descriptors developed by Cohen (7) and extended by Rosenthal (10), which are summarized in Table S1, were used for qualitative interpretation of effect sizes.

**Table S1.** Thresholds for interpreting effect sizes. The rationale for these benchmarks was outlined by Cohen (7). Supplementing Cohen’s definition of small, medium and large effect sizes, Rosenthal extended the classification to very large effects (10). *d*, standardized mean difference or effect size.

| **EFFECT SIZE** | **INTERPRETATION** |
| --- | --- |
| $0.00 < d < 0.20$ | Negligible or Trivial |
| $0.20 \leq d < 0.50$ | Small |
| $0.50 \leq d < 0.80$ | Medium |
| $0.80 \leq d < 1.30$ | Large |
| $d \geq 1.30$ | Very Large |

**Figure S1.**
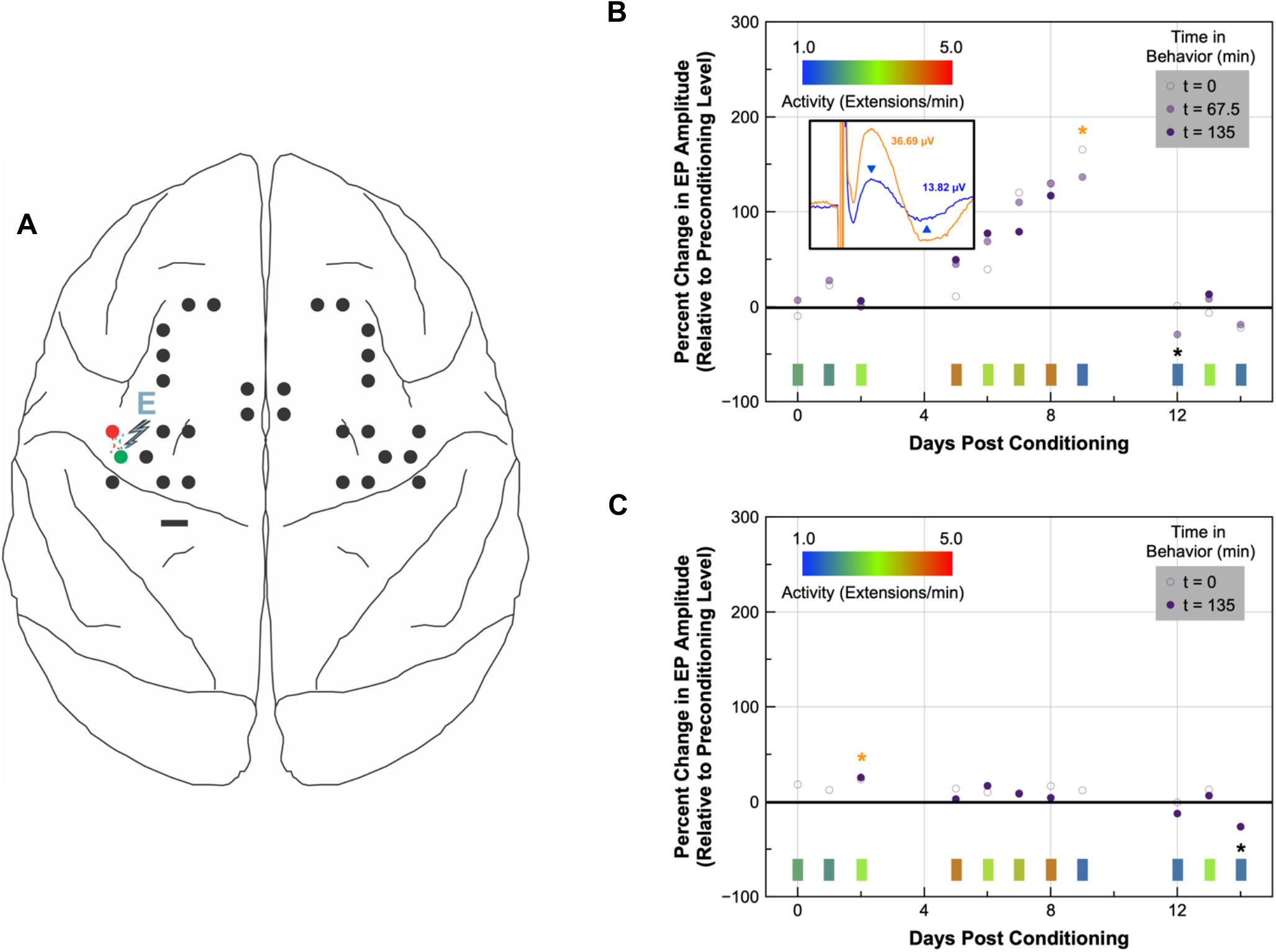
Cortical strengthening produced by combining conditioning and behavior is direction specific. (A) Schematic of the implant in monkey Y with the conditioning site pair, whose data are shown in (B) and (C), indicated in red and green. Motor outputs were assessed with preconditioning ICMS trains whose delivery at Ntest (red site) and Nstim (green site) produced activation of both wrist flexors and extensors. Extension-gated stimulation was delivered at Nstim during conditioning. E, extension; scale bar, 3 mm. (B) Test stimuli were delivered on day 0 and during behavioral sessions (at three time points; 0, 67.5, 135) that occurred every weekday over the subsequent two weeks. The percent change in EP amplitude (relative to the preconditioning level) was used to document connectivity changes in the forward direction (*i.e*., from Ntest to Nstim) during the 15-day experiment. Inset on the left shows the preconditioning EP (blue) and the EP evoked at the maximum gain (orange), with color-matched labels indicating the respective EP amplitudes. The EP amplitude was characterized as the size of the peak-to-trough (indicated by blue arrowheads) deflection, shown here for the preconditioning EP. The same peak and trough were used to characterize postconditioning EP amplitudes. (C) Documentation of connectivity changes in the reverse direction (*i.e*., from Nstim to Ntest) during the same 15-day experiment. Test stimuli were delivered at two time points (0, 135) during behavioral sessions to reduce the total amount of stimulation at Nstim (which also received the conditioning stimuli). Activity-inset and color-bar descriptions for both plots are provided in Figure 3C legend. Maxima and minima of the two datasets are indicated by orange and black asterisks, respectively.

**Figure S2.**
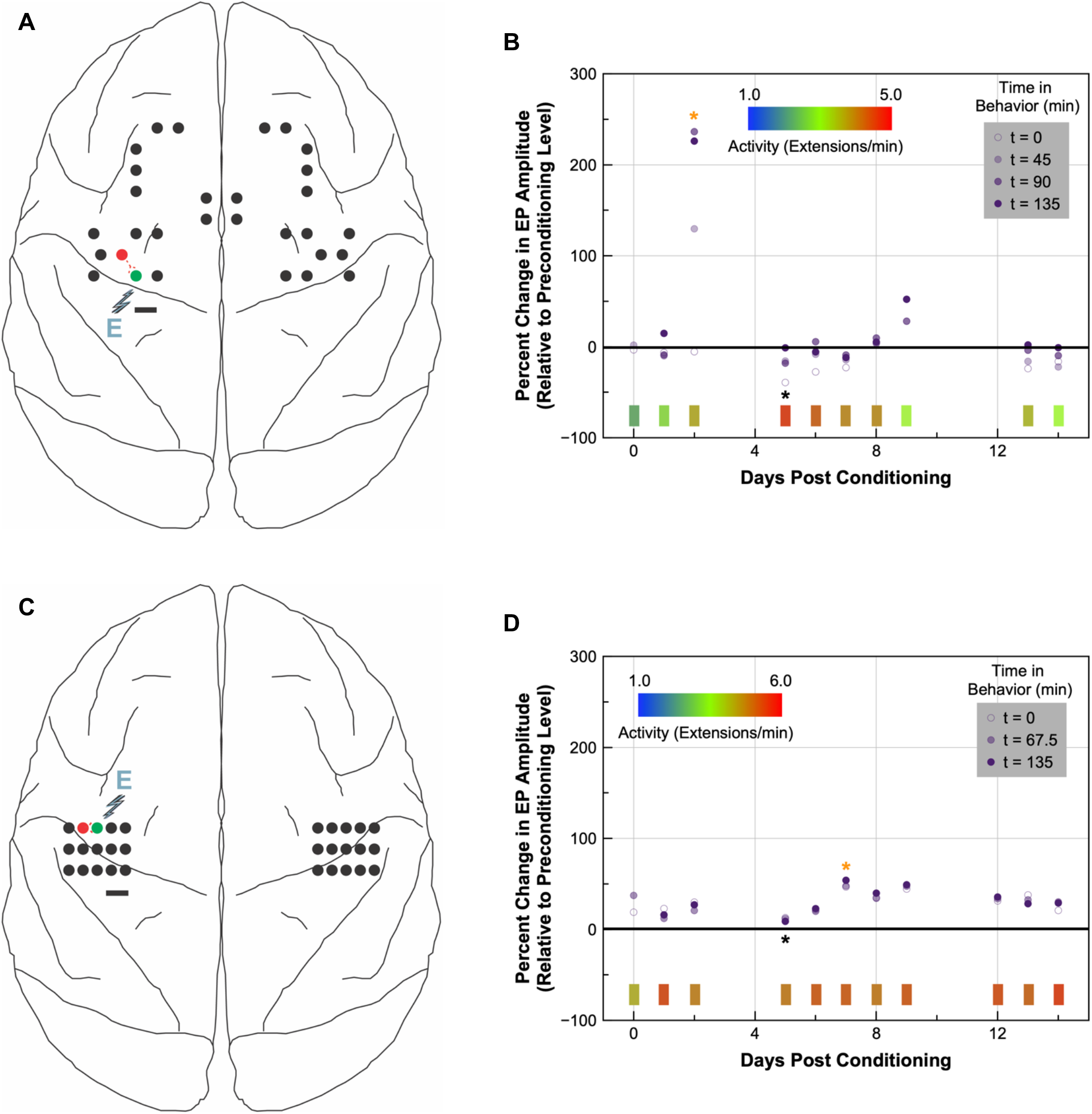
Variabilities in the magnitude and duration of cortical plasticity induced by combining conditioning and behavior. (B, D) The percent change in EP amplitude was used to document cortical-connectivity changes in the forward direction, *i.e*., from Ntest (red site) to Nstim (green site), produced by conditioning and behavior, delivered together, at two different left motor-cortical site pairs, shown in (A) and (C), respectively. Test stimuli were delivered on day 0 and during behavioral sessions, at three (0, 67.5, 135; D) or four (0, 45, 90, 135; B) time points, that occurred every weekday over the subsequent two weeks. Activity-inset and color-bar descriptions for both plots are provided in Figure 3C legend. Maxima and minima of the two datasets are indicated by orange and black asterisks, respectively. (B) and (D) show data from monkey Y and monkey U, respectively. E, extension; scale bars in (A) and (C), 3 mm.

**Figure S3.**
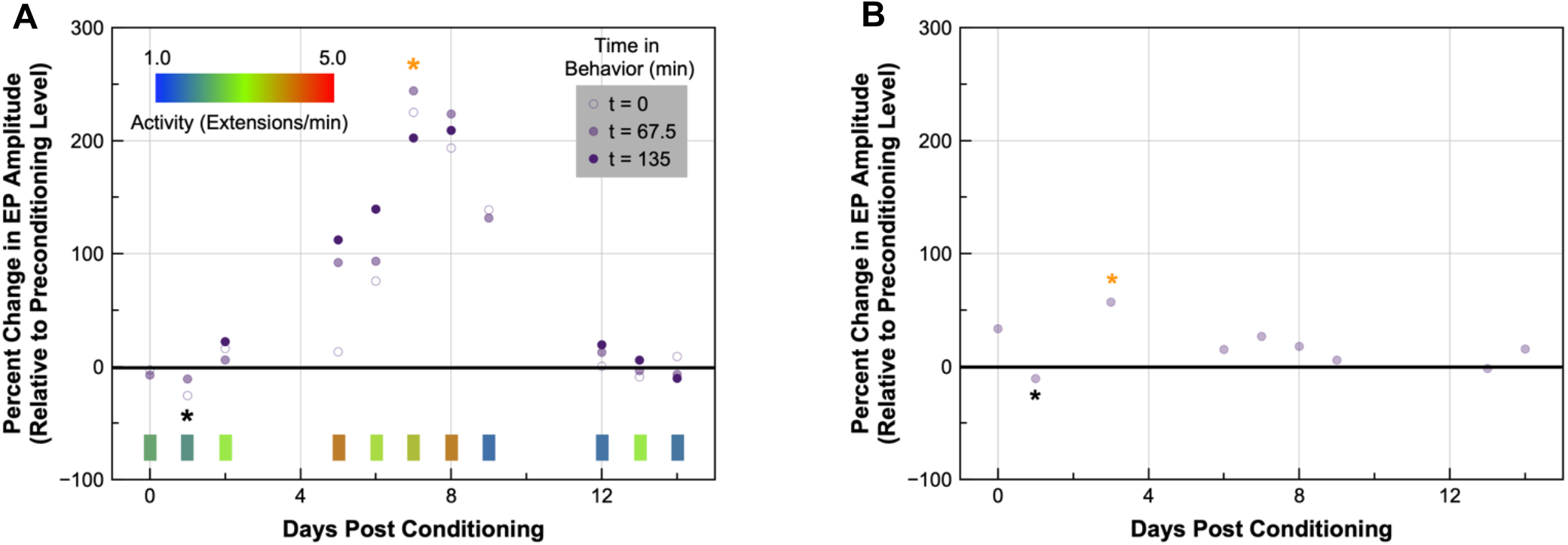
Reproducibility of effects. The percent change in EP amplitude was used to re-document cortical-connectivity changes in the forward direction produced by conditioning-and-behavior (A) and conditioning alone (B) at the same pair of left motor-cortical sites shown in Figure 3B to assess reproducibility. Test stimuli were delivered on day 0 and during following sessions, at three time points (0, 67.5, 135; A) or just at the start of each session (B), that occurred every weekday over the subsequent two weeks. Activity-inset and color-bar descriptions for (A) are provided in Figure 3C legend. Maxima and minima of the two datasets are indicated by orange and black asterisks, respectively.

**Figure S4.**
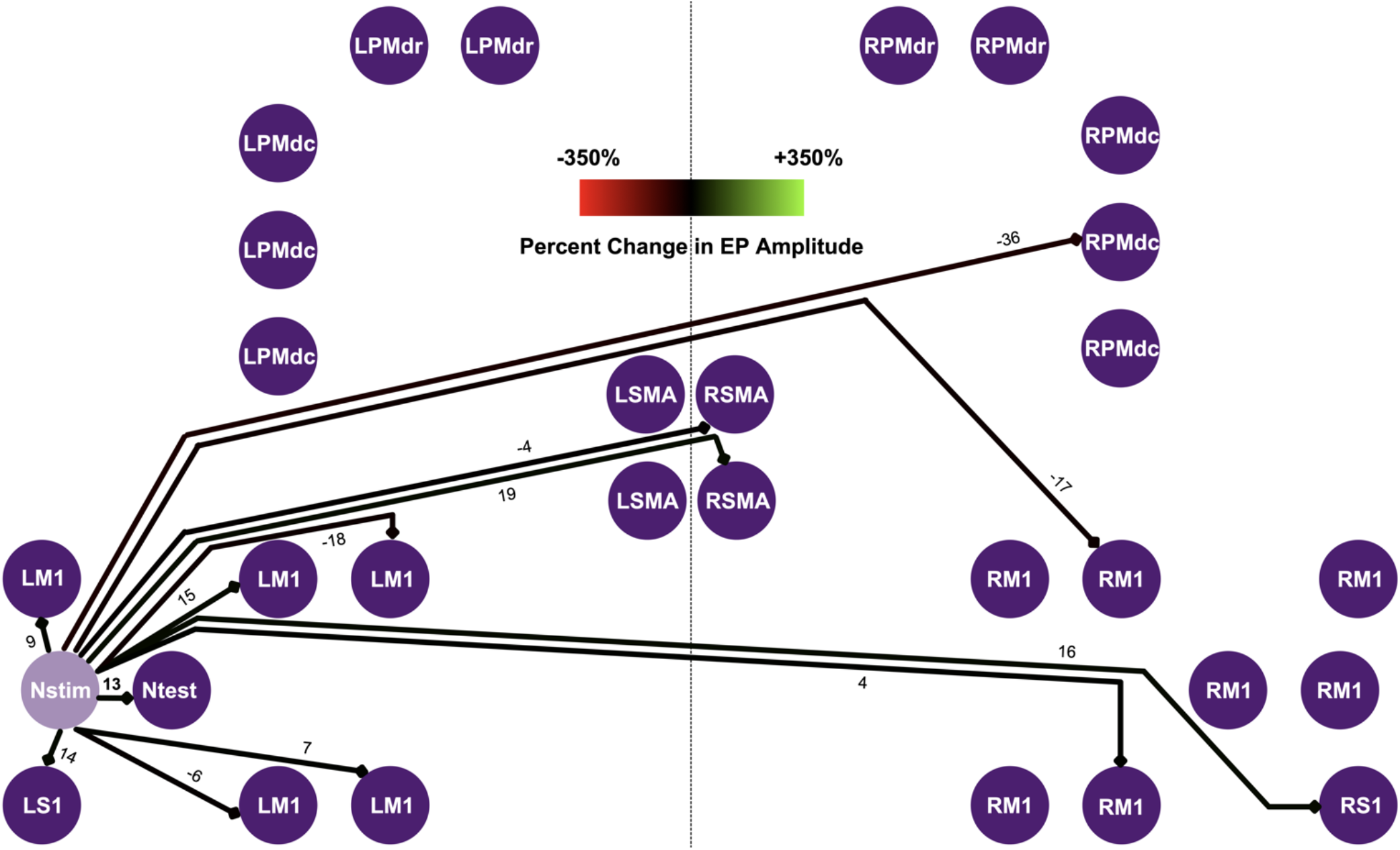
Lack of potentiation in outgoing connections from the postsynaptic site. The percent change in EP amplitude was used to document global connectivity changes produced by combining conditioning and behavior. Outgoing connections from Nstim showed little to no change when a 225% gain (Figure S3A, day 7/t = 0) was observed in the strength of the connection from Ntest to Nstim. The corresponding change from Nstim to Ntest was 13% (shown here in bold). Colored arrows denote the magnitude of connectivity changes imposed on the same graded three-color (red-black-green) scale (inset on top) used in Figure 6A; numbers indicate percent changes. Data for this figure come from the same pair of left motor-cortical conditioning sites shown in Figure 6. All other description is same as in Figure 6.

**Figure S5.**
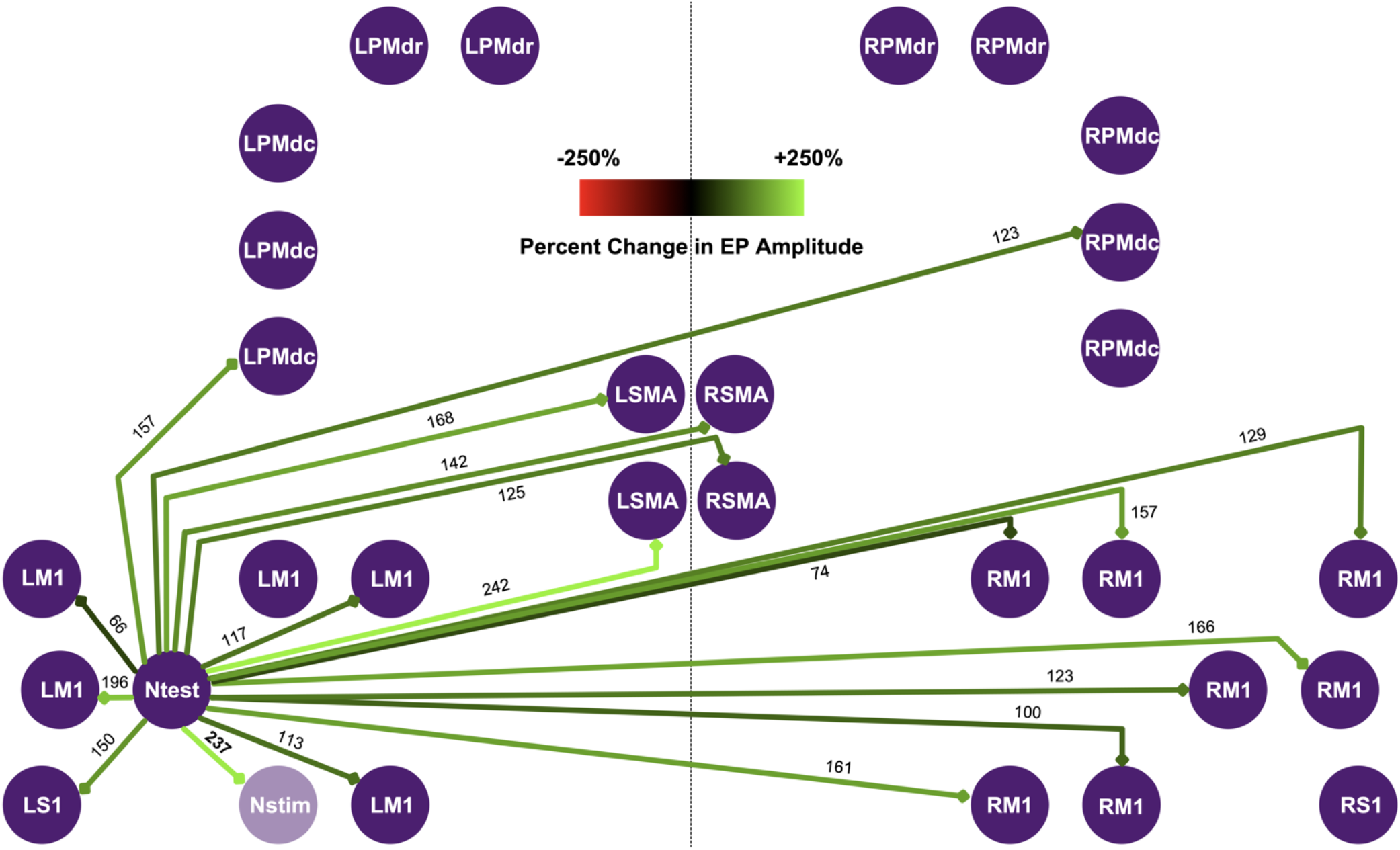
Global effects of combining conditioning and behavior. The percent change in EP amplitude was used to document global connectivity changes produced by combining conditioning and behavior. Concurrent potentiation in the strength of all outgoing connections from the presynaptic site (Ntest) was observed when there was a 237% strengthening (indicated in bold; also see orange asterisk in Figure S2B) of the connection from Ntest to Nstim. Colored arrows denote the magnitude of connectivity changes imposed on a graded three-color scale (inset on top); numbers indicate percent changes. Schematic represents the implant of monkey Y. Ntest and Nstim are sites in LM1. All other description is same as in Figure 6.

**Figure S6.**
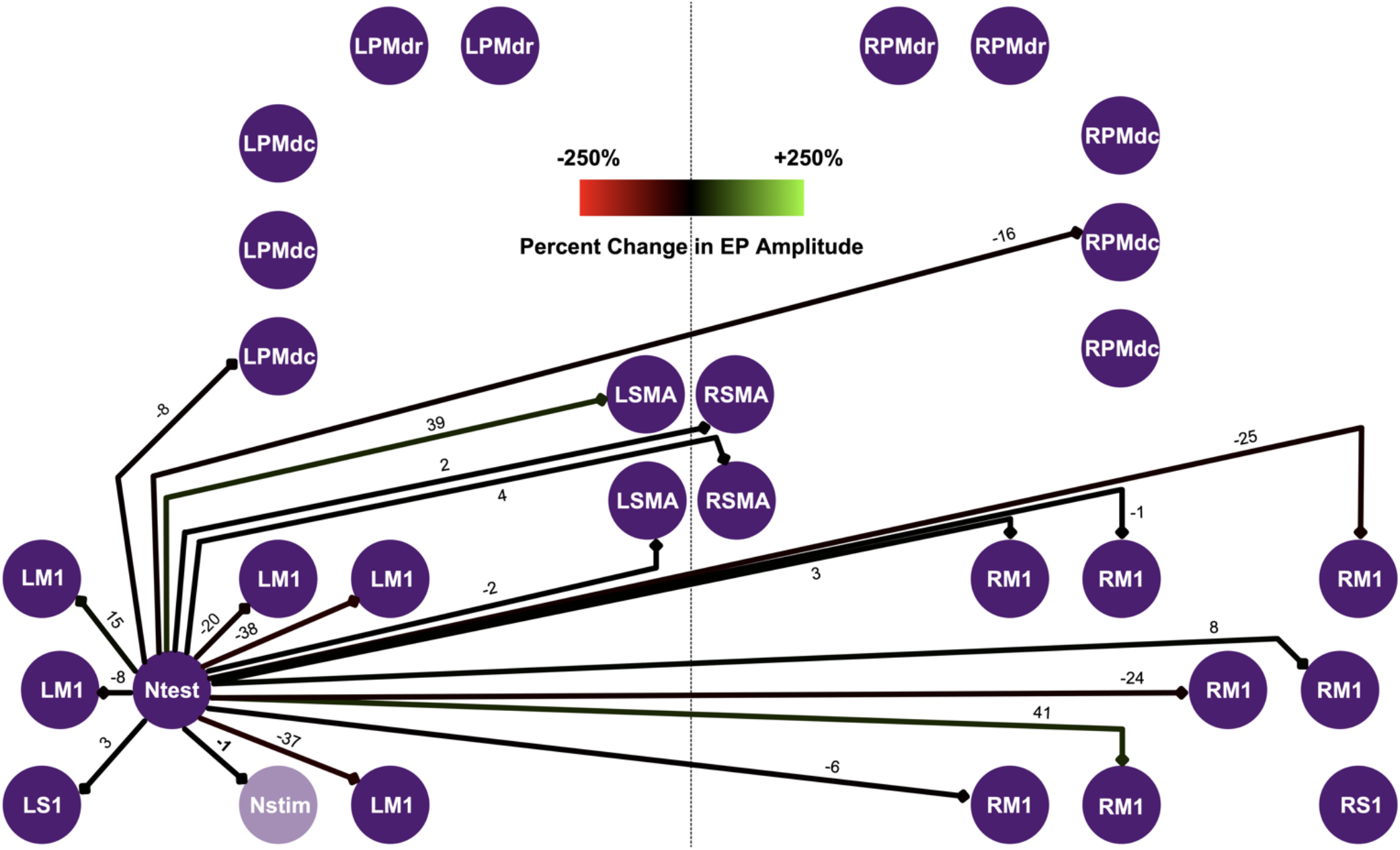
Global strengthening requires the induction of potentiation. The percent change in EP amplitude was used to document global connectivity changes produced by combining conditioning and behavior. Little to no changes in the strength of outgoing connections from Ntest were observed when there was a −1% change (indicated in bold) in the strength of the connection from Ntest to Nstim. Colored arrows denote the magnitude of connectivity changes imposed on the same graded three-color scale (inset on top) used in Figure S5; numbers indicate percent changes. Schematic represents the implant of monkey Y. Data for this figure come from the same pair of left motor-cortical conditioning sites shown in Figure S5. Implant description is same as in Figure 6.

